# Structural Disconnection of the Tool Use Network After Left Hemisphere Stroke Predicts Limb Apraxia Severity

**DOI:** 10.1101/2020.03.25.003517

**Authors:** Frank E. Garcea, Clint Greene, Scott T. Grafton, Laurel J. Buxbaum

**Author notes:** Corresponding Author Frank E. Garcea, Moss Rehabilitation Research Institute, Elkins Park, PA 19027.

## Abstract

Producing a tool use gesture is a complex process drawing upon the integration of stored knowledge of tools and their associated actions with sensory-motor mechanisms supporting the planning and control of hand and arm actions. Understanding how sensory-motor systems in parietal cortex interface with semantic representations of actions and objects in the temporal lobe remains a critical issue, and is hypothesized to be a key determinant of the severity of limb apraxia, a deficit in producing skilled action after left hemisphere stroke. We used voxel-based and connectome-based lesion symptom mapping with data from 57 left hemisphere stroke participants to assess the lesion sites and structural disconnection patterns associated with poor tool use gesturing. We found that structural disconnection between the left inferior parietal lobule, lateral temporal lobe (left middle temporal gyrus) and ventral temporal cortex (left medial fusiform gyrus) predicted the severity of tool use gesturing performance. Control analyses demonstrated that reductions in right-hand grip strength were associated with motor system disconnection, bypassing regions supporting tool use gesturing. Our findings provide causal evidence that limb apraxia may arise, in part, from disconnection of conceptual representations in the temporal lobe from mechanisms enabling skilled action production in the inferior parietal lobule.

## Introduction

The ability to produce skilled object-directed action in order to satisfy behavioral goals is a cornerstone of high-level cognitive and motor function, referred to as praxis. A central focus in the study of human praxis function is to elucidate the cognitive and neural mechanisms that interface visual processing, semantic memory, and skilled action production in order to recognize and manipulate objects in a functionally appropriate manner. A point of entry into the study of object-directed action derives from neuropsychological assessments of patients with upper limb apraxia (hereafter referred to as apraxia). Apraxia is a deficit in producing and imitating gestures that is not reducible to low-level sensory or motor dysfunction (for review, see Johnson-Frey, 2004). Patients with apraxia are impaired in producing skilled actions, often misshaping the fingers and hands when instructed to pantomime the use of objects or imitate novel gestures. Several ‘box and arrow’ information processing models (e.g., see Rothi et al., 1991; Roy & Square, 1985; Rumiati et al., 2010) posit a number of critical cognitive components that lead to different types of apraxia when lesioned or disconnected. It is only recently, however, that investigators have begun to consider how distributed cognitive models of praxis may be mapped to distributed neuroanatomic substrates (e.g., Watson et al., 2019). Understanding the consequence of specific lesion loci and disconnectivity for patterns of apraxia has implications for understanding the representation of action and object knowledge in the human brain, and provides a basis with which to identify the underlying structural connectivity interfacing object knowledge with action production processes to support skilled object-directed action.

Most cognitive models of praxis posit a distinction between a direct and indirect route for action processing (Buxbaum, 2001; Cubelli et al., 2000; Rothi et al., 1991; Rumiati & Humphreys, 1998). The direct route transforms current visual input into motor output, driven ‘bottom-up’ by the stimulus. The indirect route supports gesture production by interfacing current visual input with stored action representations. The indirect route provides a processing advantage when generating a meaningful gesture (e.g., hammering a nail), because the implementation of that gesture is facilitated by semantic memory, including the retrieval of the visual appearance of a gesture (e.g., that a swung hammer moves in a particular trajectory), function knowledge retrieval (e.g., that the function of the act of hammering is to pound in nails), and the retrieval of postural knowledge (e.g., that the configuration of the hand and fingers are positioned in a non-arbitrary manner in order to functionally grasp and manipulate a hammer; e.g., see Bracci & Peelen, 2013). In this regard, performance when gesturing tool use is a sensitive assay of the contributions of the indirect route, as the visual input must interface with the semantic system for accurate demonstration of tool use. By contrast, imitation of meaningless gestures provides a means with which to evaluate the integrity of the direct route, because the transformation of visual input into sensory-motor plans occurs without access to semantic information (for review, see Binkofski & Buxbaum, 2013).

The left inferior parietal lobule is a core region supporting praxis function, as lesions to this region (supramarginal gyrus, angular gyrus) are associated with spatial and temporal errors when apraxics pantomime or imitate actions (Garcea et al., 2013; Goldenberg & Spatt, 2009; Mengotti et al., 2013; Negri et al., 2007; for review, see Buxbaum & Kalenine, 2010; Goldenberg, 2009; Leiguarda & Marsden, 2000; Rumiati et al., 2010). It is noteworthy that imitation of meaningless gesture is more strongly associated with lesions to the left inferior parietal lobule (Hoeren et al., 2014; Tessari et al., 2007; Weiss et al., 2001; for a neuroimaging meta-analysis, see Caspers et al., 2010), whereas tool use gesturing draws on additional stored representations, including visual and postural knowledge of actions that are processed in the posterior temporal lobe. For example, although spatiotemporal errors in meaningless gesture imitation are associated with parietal and pre-motor lesions, hand posture errors in tool use gesturing, reflecting knowledge of the non-arbitrary hand and finger configuration required for functional manipulation, are associated with lesions to the left posterior middle temporal gyrus (Buxbaum et al., 2014; see also Hoeren et al., 2014; Manuel et al., 2013).

Neuropsychological data provide evidence that the left posterior middle temporal gyrus supports action recognition and tool knowledge, as lesions to this region are associated with impairment when matching videos of tool use gestures to action verbs (Kalenine et al., 2010; Kemmerer et al., 2012) and impaired performance when naming tools but not animals (Brambati et al., 2006; Campanella et al., 2010). Functional magnetic resonance imaging (fMRI) findings in neurotypical adults are consistent with the available neuropsychological data, as there is increased blood oxygen level-dependent (BOLD) contrast in the left posterior middle temporal gyrus when participants gesture tool use (Brandi et al., 2014; Johnson-Frey et al., 2005; Vry et al., 2015), view images of tools (Beauchamp et al., 2002; Chao et al., 1999; Garcea et al., 2016; Mahon et al., 2007; for review, see Lingnau & Downing, 2015; A. Martin, 2007), and make judgments about actions (Kable et al., 2005; Kable et al., 2002; Wurm & Caramazza, 2019), including tool use actions (Kleineberg et al., 2018; Valyear & Culham, 2010). Furthermore, an emerging literature demonstrates there is increased functional connectivity between the left inferior parietal lobule and left posterior middle temporal gyrus when neurotypical participants gesture the use of tools (Garcea et al., 2018; Hutchison & Gallivan, 2018; Vingerhoets & Clauwaert, 2015). Consistent with this finding, the degree of resting state functional connectivity reduction between the left inferior parietal lobule and left posterior middle temporal gyrus predicts the severity of tool use gesturing deficits in participants with apraxia after left cerebrovascular accident (LCVA) (Watson et al., 2019).

The left medial fusiform gyrus, a region implicated in fMRI studies of tool processing (Chao et al., 1999; Garcea & Mahon, 2014; Mahon et al., 2007), is sensitive to material and textural properties of manipulable objects (Cant et al., 2009; Cant & Goodale, 2007), and is another region for which disconnection appears related to praxis capacity. Tool use gesturing deficits in apraxia are associated with reduced resting state functional connectivity between the inferior parietal lobule and the left medial fusiform gyrus (Watson et al., 2019). Moreover, in pre-operative neurosurgery participants the degree of reduced BOLD contrast for tools in the left medial fusiform gyrus is associated with lesions in the left inferior parietal lobule (Garcea, Almeida, et al., 2019), suggesting that parietal lesions disrupt the processing of tools in functionally connected nodes in ventral temporal cortex.

Other nodes of relevance to praxis are pre-central and prefrontal cortices, including left ventral premotor cortex and left inferior and middle frontal gyri. Lesions to the left inferior frontal or middle frontal gyri can result in tool gesturing deficits (Goldenberg et al., 2007; Haaland et al., 2000; see also Bohlhalter et al., 2011), and both regions exhibit increased functional connectivity to the left inferior parietal lobule when neurotypical participants gesture tool use (Garcea & Buxbaum, 2019). In addition, greater BOLD contrast is observed in the left ventral premotor cortex when participants judge the appropriateness of an action deployed to an object, as this activation is driven by the number of potential actions associated with the object (Schubotz et al., 2014; for review, see Buxbaum, 2017).

In sum, these findings indicate that in the course of generating a tool use action, conceptual attributes of actions (e.g., knowledge of hand posture and action function) and visual attributes of objects (e.g., object form, surface texture) provide key inputs to parietal sensory-motor systems in order to manipulate a tool skillfully and in accordance with its function. These action and object processes, by hypothesis, interface with frontal-motor regions critical for action selection and motor specification. We (and others) refer to the network of brain regions that collectively support the ability to recognize actions and use manipulable objects as the Tool Use Network (see Buxbaum & Randerath, 2018; Garcea & Mahon, 2014). Recent findings suggest that unique variance in apraxia severity may be captured by disruption of functional connectivity among nodes in the Tool Use Network (Watson et al., 2019); however, given the paucity of structural connectivity research in apraxia, the goal of this study was to investigate the extent to which structural disconnection among nodes of the Tool Use Network is predictive of deficits in tool use gesturing.

Recent studies have raised the concern that stroke lesions invading white matter can distort the quality of diffusion data, reducing the accuracy of whole-brain tractography quantification (de Groot et al., 2013; Langen et al., 2018; Theaud et al., 2017). One proposal to overcome this limitation is to compute a participant-specific (dis)connectome on the basis of inferred structural disconnection given a participant’s lesion location in relation to structural connectivity in a normative dataset. Greene and colleagues (2019) recently developed an analytic pipeline that infers structural disconnection from lesion location, and demonstrated that the extent of disconnection of the corticospinal tract induced by stroke lesions could reliably predict grip strength scores over and above lesion volume, owing to the fact small lesions can have disruptive effects if they encroach upon white matter tracts. In the current project, we used Greene and colleagues’ analytic pipeline to investigate structural disconnection associated with deficits in tool use gesturing.

Fifty-seven LCVA participants took part in neuropsychological testing of tool use gesturing and meaningless gesture imitation using the ipsilesional hand, and underwent high-resolution structural neuroimaging. We then used support vector machines in combination with voxel-based and connectome-based lesion symptom mapping to investigate the lesion sites and disconnection patterns, respectively, associated with reduced tool use gesturing performance, controlling for performance in meaningless gesture imitation. We reasoned that removing variance in tool use gesturing shared with meaningless gesture imitation will, by hypothesis, permit a test of structural disconnection associated with impaired retrieval of stored knowledge of tools and their associated actions, which would not be present in the context of imitating a novel, meaningless gesture.

In a second aim, we investigated lesion sites and disconnection patterns associated with motor system disconnection, as 41 of the 57 participants also took part in a test of grip strength using the ipsilesional and contralesional hand. We reasoned that the pattern of disconnection among nodes in the Tool Use Network associated with tool use gesturing severity should not predict weakened contralesional grip strength. By contrast, we hypothesized that disconnection among pre- and post-central gyri, and subcortical structures in the basal ganglia would be predictive of reduced grip strength of the contralesional hand after LCVA.

## Materials and Methods

### Participants

Sixty-six chronic LCVA participants were recruited from the Neuro-Cognitive Research Registry at Moss Rehabilitation Research Institute. Of those participants, 58 completed the gesturing tool use task and the meaningless imitation task. One of the 58 participants was determined to be an outlier, and thus all final analyses included 57 participants (28 female; mean age = 57.5 years, SD = 11.6 years, range = 31 – 80 years; mean education = 14.2 years, SD = 2.7 years, range = 9 – 21 years). Participants were all right-hand dominant (1 reported as ambidextrous) and had suffered a single left hemisphere stroke at least 3 months prior to testing (mean number of months since stroke = 40 months, SD = 46.4 months; range = 4 – 184 months). Participants were excluded if they had a history of psychosis, drug or alcohol abuse, co-morbid neurological disorder, or severe language comprehension deficits established with the Western Aphasia Battery (Kertesz, 1982). See Table 1 for demographic information and Figure 1 for cortical and subcortical lesion distribution for each LCVA participant.

**Table 1.** Demographic information and lesion volume for each LCVA participant. Underscores indicate participants for whom measures of right- and left-hand grip strength were not available. *Abbreviations*. GTS, Gesturing tool use to the sight of objects; IMI, meaningless imitation; HP, hand posture accuracy; LH, left hand; RH, right hand

| LCVA Participant | GTS HP (%) | IMI HP (%) | LH Grip Strength | RH Grip Strength | Months Post Stroke | Lesion Volume (1mm<sup>3</sup>) | Years of Education | Gender | Age |
| --- | --- | --- | --- | --- | --- | --- | --- | --- | --- |
| 1 | 0.60 | 0.70 | 22.34 | 20.34 | 174 | 115118 | 16 | F | 54 |
| 2 | 0.75 | 0.50 | — | — | 172 | 258736 | 18 | M | 68 |
| 3 | 1.0 | 0.90 | — | — | 7 | 16978 | 12 | M | 31 |
| 4 | 0.83 | 0.60 | 37.67 | 27 | 112 | 166393 | 13 | M | 57 |
| 5 | 0.64 | 0.60 | 14.67 | 15 | 7 | 20190 | 12 | F | 69 |
| 6 | 0.73 | 0.70 | 30 | 20 | 14 | 76301 | 18 | M | 64 |
| 7 | 0.55 | 0 | — | — | 22 | 171128 | 18 | M | 51 |
| 8 | 0.85 | 0.90 | 30.67 | 33 | 89 | 51780 | 21 | M | 79 |
| 9 | 0.83 | 0.60 | 20.67 | 16 | 47 | 82964 | 14 | M | 53 |
| 10 | 0.60 | 0.70 | 15 | 14 | 4 | 33183 | 14 | F | 59 |
| 11 | 0.93 | 0.70 | 17 | 13 | 73 | 37091 | 14 | F | 55 |
| 12 | 0.54 | 0.24 | 13.34 | 15.34 | 114 | 16547 | 12 | F | 80 |
| 13 | 0.83 | 0.50 | — | — | 84 | 80020 | 13 | F | 47 |
| 14 | 0.88 | 0.70 | 47.67 | 37.34 | 24 | 47442 | 12 | M | 53 |
| 15 | 0.62 | 0.60 | — | — | 31 | 179606 | 15 | M | 57 |
| 16 | 0.68 | 0.70 | 31 | 33 | 6 | 23141 | 12 | M | 55 |
| 17 | 1.0 | 0.60 | 37.67 | 31.67 | 53 | 20105 | 21 | M | 56 |
| 18 | 0.8 | 0.70 | — | — | 8 | 71022 | 16 | F | 60 |
| 19 | 0.57 | 0.40 | 14.67 | 22 | 9 | 62204 | 18 | F | 32 |
| 20 | 0.93 | 0.50 | 24.34 | 29.67 | 68 | 94536 | 12 | F | 45 |
| 21 | 0.25 | 0.50 | — | — | 11 | 303310 | 12 | F | 65 |
| 22 | 0.50 | 0.40 | 23.34 | 28 | 7 | 61198 | 12 | F | 71 |
| 23 | 0.66 | 0.67 | — | — | 32 | 136576 | 12 | F | 46 |
| 24 | 0.75 | 0.70 | 33.34 | 26 | 24 | 52416 | 14 | M | 48 |
| 25 | 0.84 | 0.70 | 18 | 22 | 9 | 128897 | 12 | F | 54 |
| 26 | 0.77 | 0.60 | 35.67 | 34 | 7 | 51399 | 18 | M | 62 |
| 27 | 0.77 | 0.80 | — | — | 7 | 20790 | 12 | F | 50 |
| 28 | 0.85 | 0.60 | — | — | 10 | 88046 | 13 | M | 31 |
| 29 | 0.90 | 0.70 | 33.34 | 4.34 | 65 | 27840 | 12 | F | 39 |
| 30 | 0.83 | 0.50 | 29 | 16 | 6 | 92744 | 12 | M | 48 |
| 31 | 0.70 | 0.70 | — | — | 31 | 200079 | 12 | M | 61 |
| 32 | 0.64 | 0.50 | 26.67 | 8.67 | 19 | 117809 | 12 | F | 41 |
| 33 | 0.80 | 0.40 | 22 | 24.67 | 40 | 5953 | 16 | M | 64 |
| 34 | 0.70 | 0.30 | 22.67 | 22.67 | 9 | 32684 | 13 | F | 64 |
| 35 | 0.95 | 0.40 | 46.34 | 54 | 32 | 337 | 16 | M | 55 |
| 36 | 0.55 | 0.90 | 28 | 30.34 | 31 | 32003 | 18 | M | 71 |
| 37 | 0.95 | 0.60 | 36 | 14.34 | 23 | 4095 | 18 | M | 63 |
| 38 | 0.87 | 0.70 | 42.34 | 14 | 10 | 27004 | 12 | M | 48 |
| 39 | 0.77 | 0.90 | 26.34 | 34 | 20 | 11961 | 12 | M | 58 |
| 40 | 0.87 | 0.40 | 31.34 | 35.67 | 17 | 102522 | 16 | F | 38 |
| 41 | 0.90 | 0.70 | 34.34 | 22.34 | 16 | 14660 | 18 | M | 53 |
| 42 | 0.80 | 0.70 | 21.67 | 20 | 4 | 9489 | 12 | F | 68 |
| 43 | 0.85 | 0.40 | 35.67 | 0 | 63 | 26714 | 18 | F | 46 |
| 44 | 0.90 | 0.90 | 19.67 | 17.67 | 17 | 16977 | 9 | M | 50 |
| 45 | 0.90 | 0.80 | 23.67 | 17.34 | 6 | 30414 | 12 | F | 68 |
| 46 | 0.66 | 0.60 | — | — | 7 | 16186 | 14 | F | 70 |
| 47 | 0.98 | 1.0 | 46 | 53.34 | 6 | 64375 | 12 | M | 45 |
| 48 | 0.75 | 0.60 | 26.67 | 0 | 9 | 56281 | 9 | F | 66 |
| 49 | 0.40 | 0.40 | — | — | 12 | 225021 | 16 | M | 64 |
| 50 | 0.88 | 0.72 | 26 | 7.67 | 71 | 48459 | 13 | F | 53 |
| 51 | 0.91 | 0.86 | — | — | 17 | 93628 | 16 | F | 37 |
| 52 | 0.82 | 0.90 | 21.67 | 27.67 | 12 | 1869 | 12 | F | 67 |
| 53 | 0.98 | 0.80 | 36.67 | 32.34 | 12 | 17706 | 12 | M | 51 |
| 54 | 0.90 | 0.69 | 37.34 | 33.34 | 151 | 89072 | 16 | F | 55 |
| 55 | 0.85 | 0.72 | — | — | 184 | 62530 | 13 | F | 60 |
| 56 | 0.87 | 0.65 | 23.67 | 10.00 | 30 | 87120 | 18 | M | 76 |
| 57 | 0.80 | 0.58 | — | — | 161 | 96196 | 13 | M | 60 |

**Figure 1.**
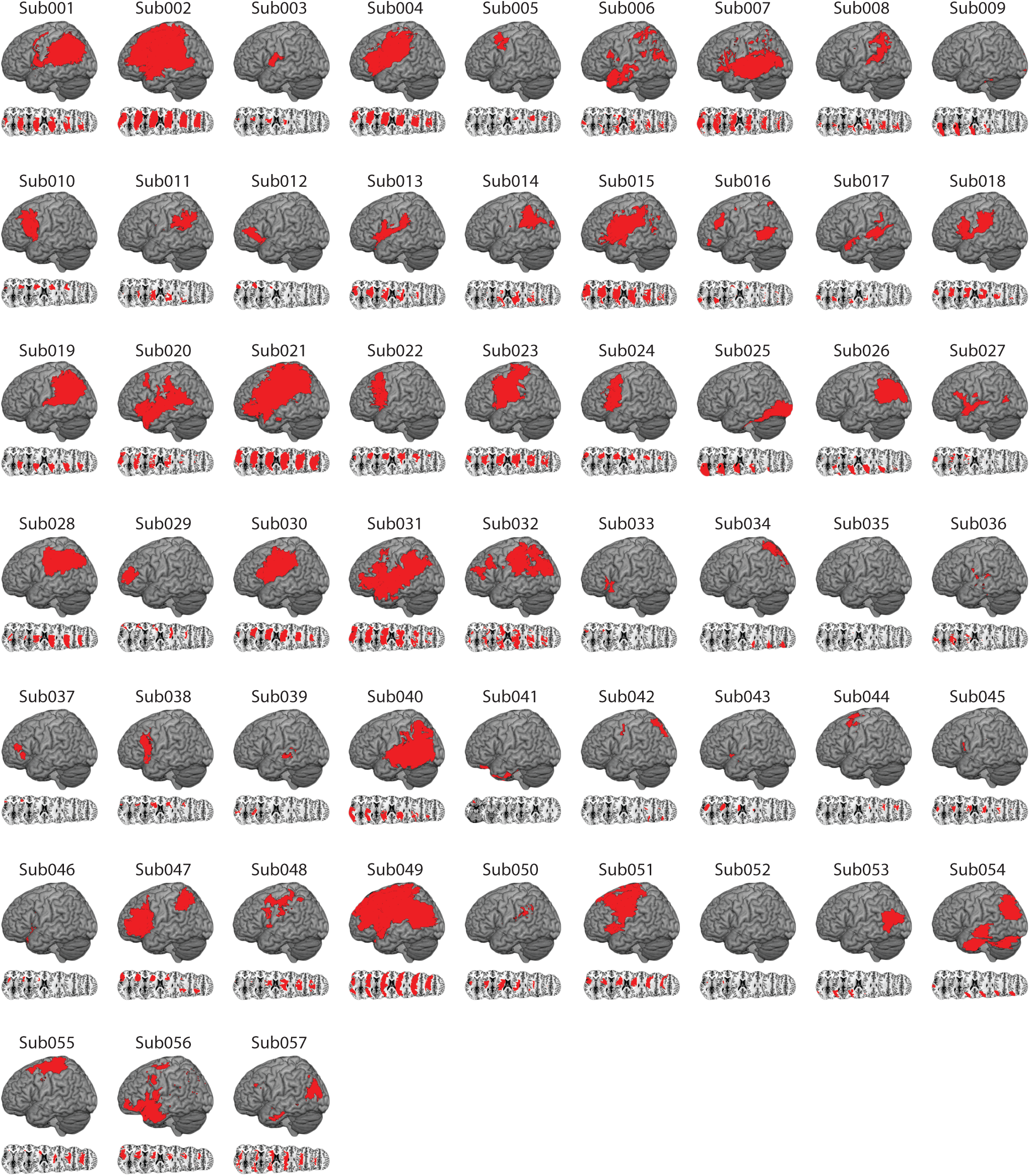
Distribution of Cortical and Subcortical Damage in Each LCVA Participant. A depiction of the 57 LCVA participants’ lesions are presented on the ch2bet template. Lesions are projected on the cortical surface but include subcortical voxels using a 12-voxel search depth. Axial slices begin at the MNI coordinate [0,0,0] and increase in 10-mm increments superiorly.

In compliance with the guidelines of the Institutional Review Board of Einstein Healthcare Network, all participants gave informed consent and were compensated for travel expenses and participation. The informed consents obtained did not include permission to make data publicly available. Accordingly, the conditions of our ethical approval do not permit anonymized study data to be publicly archived. To obtain access to the data, individuals should contact the corresponding author. Requests for data are assessed and approved by the Institutional Review Board of Einstein Healthcare Network.

### Neuropsychological Testing of Tool Use Gesturing and Meaningless Imitation

#### Gesturing Tool Use to the Sight of Objects

The gesturing tool use to the sight of objects test included 40 photographs of manipulable objects (tools) taken from the BOSS database (Brodeur et al., 2010). Tools included items with distinct use actions, including construction tools (e.g., wrench), household articles (e.g., teapot), office supplies (e.g., scissors), and bathroom items (e.g., razor). Each trial of the test began with the presentation of a 400-by-400 pixel color photograph of a tool on a computer monitor. Participants were asked to “show how you would use the tool as if you were holding and using it” with the left hand. Four practice trials with feedback (using items different than in the task itself) were given at the start of the task. As per Rothi et al. (1991), if a participant gestured the action as if their hand was the tool (body-part-as-object error), they were reminded to “show how you would use the tool as if you were actually holding it in your hand”. The first of these errors was corrected and the participant was permitted a second try (for precedent, see Garcea, Stoll, et al., 2019; Tarhan et al., 2015; Watson & Buxbaum, 2015).

#### Test of Meaningless Imitation

Participants were presented with videos of an experimenter performing 10 novel gestures and were instructed to imitate the gesture. Gestures were presented twice on each trial; during the first presentation, participants were instructed to watch the gesture in its entirety; at the beginning of the second presentation a sound was presented cueing participants to begin gesturing. The 10 novel gestures were developed to maintain similar motor characteristics of tool use gestures (e.g., plane of movement; joints moved; hand posture), but were designed such that the movement was meaningless (e.g., see Buxbaum et al., 2014). Eight of the 57 individuals took part in a version of the test with 14 meaningless actions to imitate, 10 of which were identical to the 10 meaningless actions that the remaining 49 participants gestured.

#### Coding of Action Data

Gesturing tool use and meaningless imitation tests were recorded with a digital camera and scored offline by two trained, reliable coders (Cohen’s Kappa score = 94%) who also demonstrated inter-rater reliability with previous coders in the Buxbaum lab (Cohen’s Kappa > 85%; see e.g., Buxbaum et al., 2005). Both tests were coded using a portion of the detailed praxis scoring guidelines used in our previous work (see Buxbaum et al., 2000; Buxbaum et al., 2005; Watson & Buxbaum, 2015). In the gesturing tool use test, each gesture was given credit for semantic content unless a participant performed a recognizable gesture appropriate for a semantically-related tool. Only gestures that were given credit for semantic content were scored on other action dimensions (e.g., spatiotemporal hand posture errors). Across tests of tool use gesturing and meaningless imitation, hand posture errors were assigned if the shape or movement trajectory of the hand and/or wrist was flagrantly incorrect, or if the hand or finger was used as part of the tool (i.e., body-part-as-object error, Buxbaum et al., 2005; Watson & Buxbaum, 2015). This allowed us to investigate hand posture errors in the context of producing meaningful and meaningless actions, and to isolate unique variance when generating errors in meaningful gesture production to test hypotheses of Tool Use Network disconnection in association with reduced performance in tool use gesturing.

Following action coding, we computed the average proportion of hand posture errors from the tool use gesturing test and from the meaningless imitation test. Consistent with past findings in our lab (Buxbaum et al., 2014), there was a significant relation between proportion of errors in tool use gesturing and meaningless imitation (r(55) = 0.42, *p* < .01). Thus, we regressed performance from the gesturing tool use test on the meaningless imitation test to obtain a residual hand posture error score; negative-going residual scores indicate worse performance when gesturing tool use relative to meaningless imitation, which was the focus of the current investigation. Residual scores were entered as the principal dependent variable in a support vector regression voxel-based lesion symptom mapping analysis (Figure 2B), and in a support vector regression connectome-based lesion symptom mapping analysis (Figure 3).

**Figure 2.**
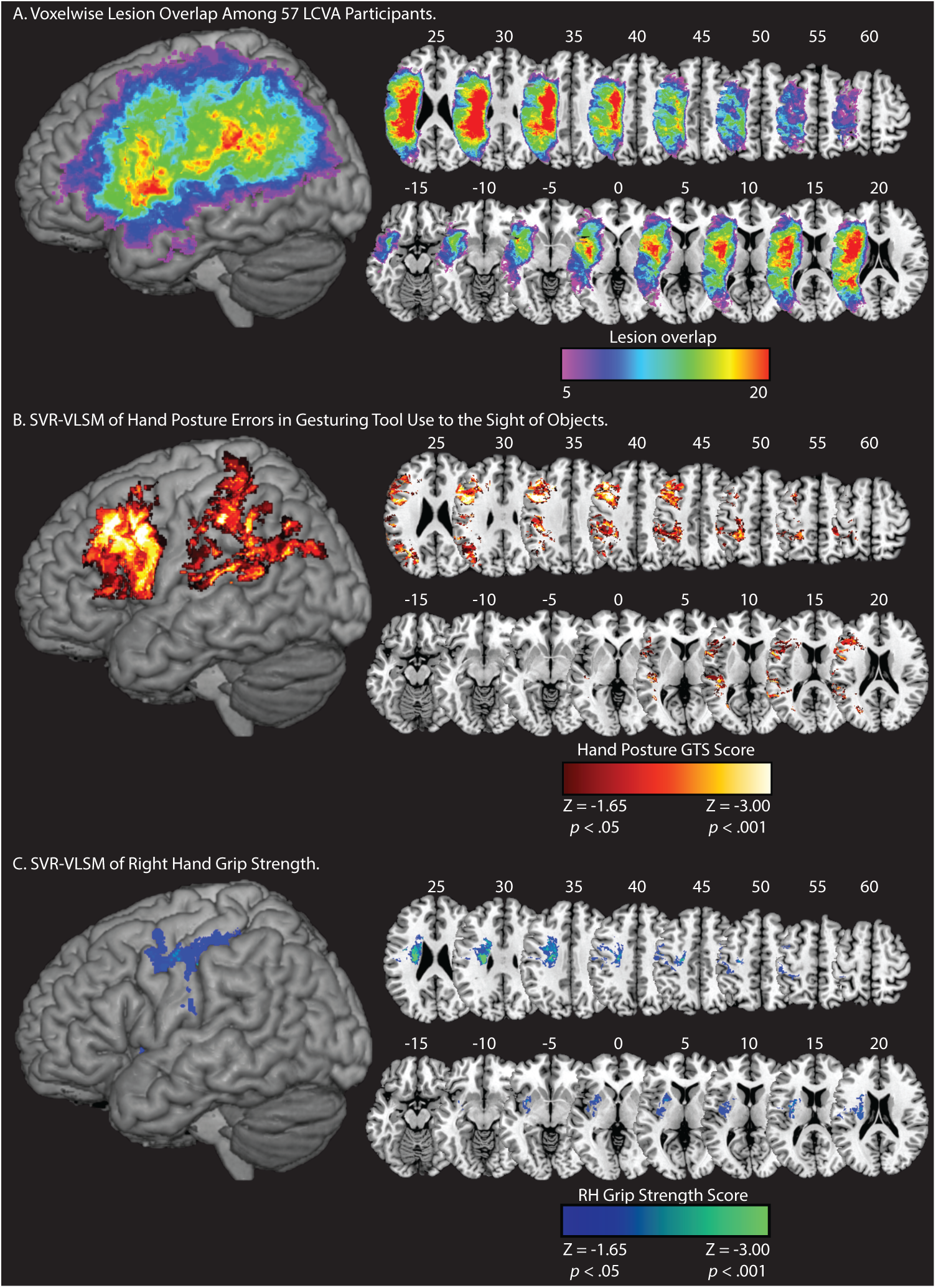
Lesion Overlap and Support Vector Regression Lesion Symptom Mapping Results. **A**. Voxelwise lesion overlap among 57 participants. Only voxels with at least 5 voxels are included in the analysis. **B**. Voxels associated with reduced performance in tool use gesturing (greater hand posture errors in tool use gesturing controlling for hand posture errors in meaningless imitation; red-to-white scale). **C**. Voxels associated with reduced right-hand grip strength (weaker grip strength with the right hand controlling for grip strength of the left hand; blue-to-green scale). Whole-brain results are rendered in MNI space in 5-mm increments. SVR-VLSM maps are set to a voxelwise threshold of *p* < .05 with 10,000 iteration Monte Carlo style permutation analysis; clusters are included if they are greater than or equal to 500 contiguous 1 mm^3^ voxels.

**Figure 3.**
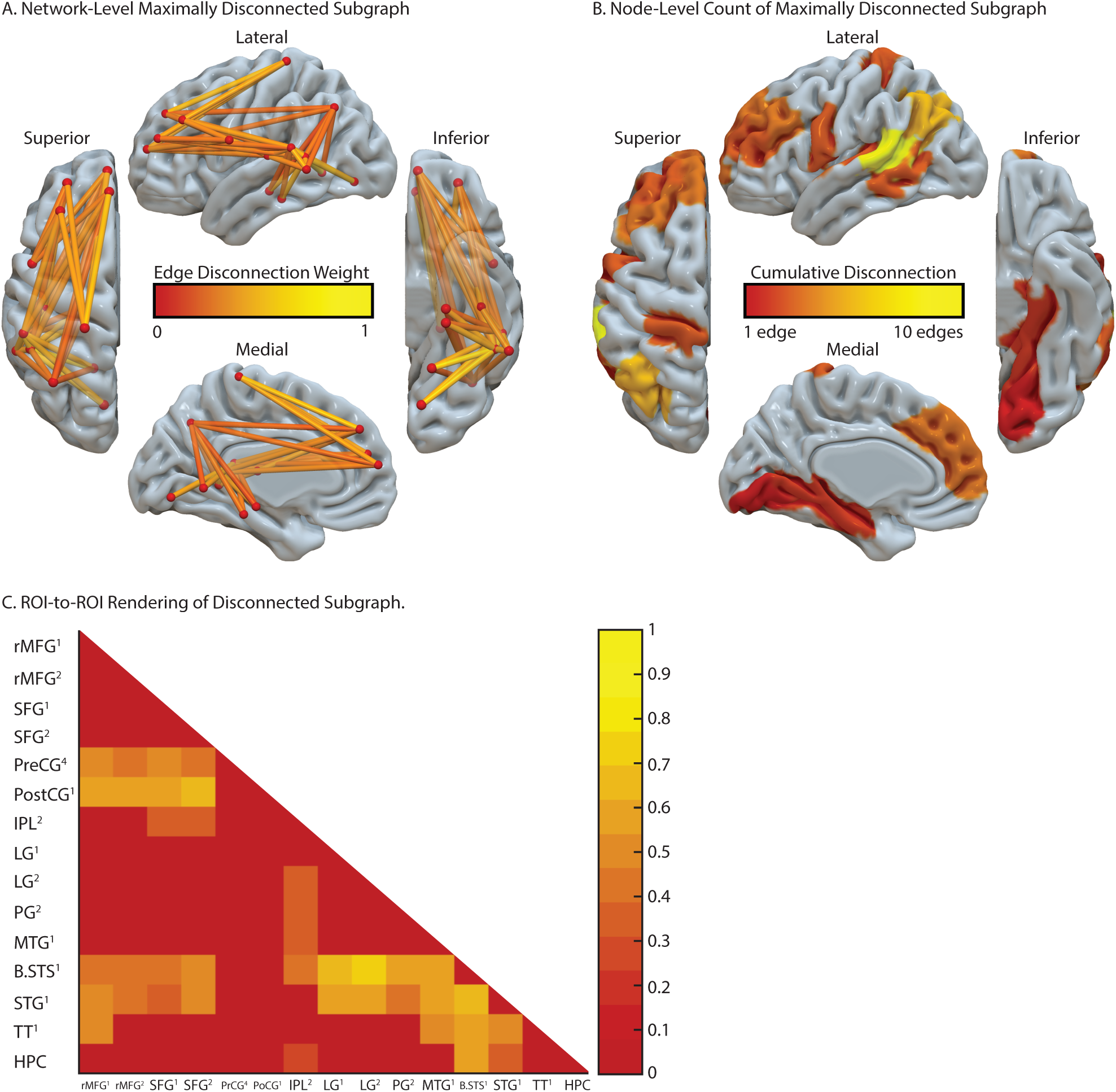
Support Vector Regression Connectome-based Lesion Symptom Mapping of Errors in Tool Use Gesturing. **A**. Maximally disconnected edges associated with reduced performance in tool use gesturing (greater hand posture errors in tool use gesturing controlling for hand posture errors in meaningless imitation and total lesion volume; red-to-yellow scale). **B**. Cumulative node-level disconnection associated with reduced tool use gesturing (red-to-yellow scale) depicted on the cortical surface. **C**. A heatmap of the disconnected subgraph depicts the ROI-to-ROI disconnection patterns projected in Fig. 3A. *Abbreviations*. rMFG, rostral middle frontal gyrus; SFG, superior frontal gyrus; PreCG, precentral gyrus; PostCG, postcentral gyrus; IPL, inferior parietal lobule; LG, lingual gyrus; PPG, parah
ippocampal gyrus; MTG, middle temporal gyrus; B. STS, bank of the superior temporal sulcus; STG, superior temporal gyrus; TT, transverse temporal. Numbers in subscript indicate anatomical subregions from the Lausanne atlas.

#### Test of Grip Strength

Forty-one of the 57 participants took part in a test of grip strength. On each trial, participants squeezed a hydraulic hand dynamometer as hard as possible using their contralesional (right) and ipsilesional (left) hand (for product details, see https://www.3bscientific.com/product-manual/W50175.pdf). Participants did not use their right hand if they had excessive weakness or poor control of the fingers or hand (likely reflecting the presence of post-stroke hemiparesis); for this reason, we did not obtain grip strength in 16 individuals. Participants took part in three trials to ensure an accurate recording of grip strength; we then averaged across the three trials to obtain a final right- and left-hand grip strength score. There was a significant correlation between right- and left-hand grip strength scores (r(39) = 0.48, *p* < .01). Thus, we regressed right-hand grip strength on left-hand grip strength to obtain a residual right-hand grip strength score; negative-going residual scores indicate reduced right-hand grip strength relative to left-hand grip strength, which was the focus of the current investigation. Residual scores were then entered as the principal dependent variable in a support vector regression voxel-based lesion symptom mapping analysis (Figure 2C), and in a support vector regression connectome-based lesion symptom mapping analysis (Figure 4).

**Figure 4.**
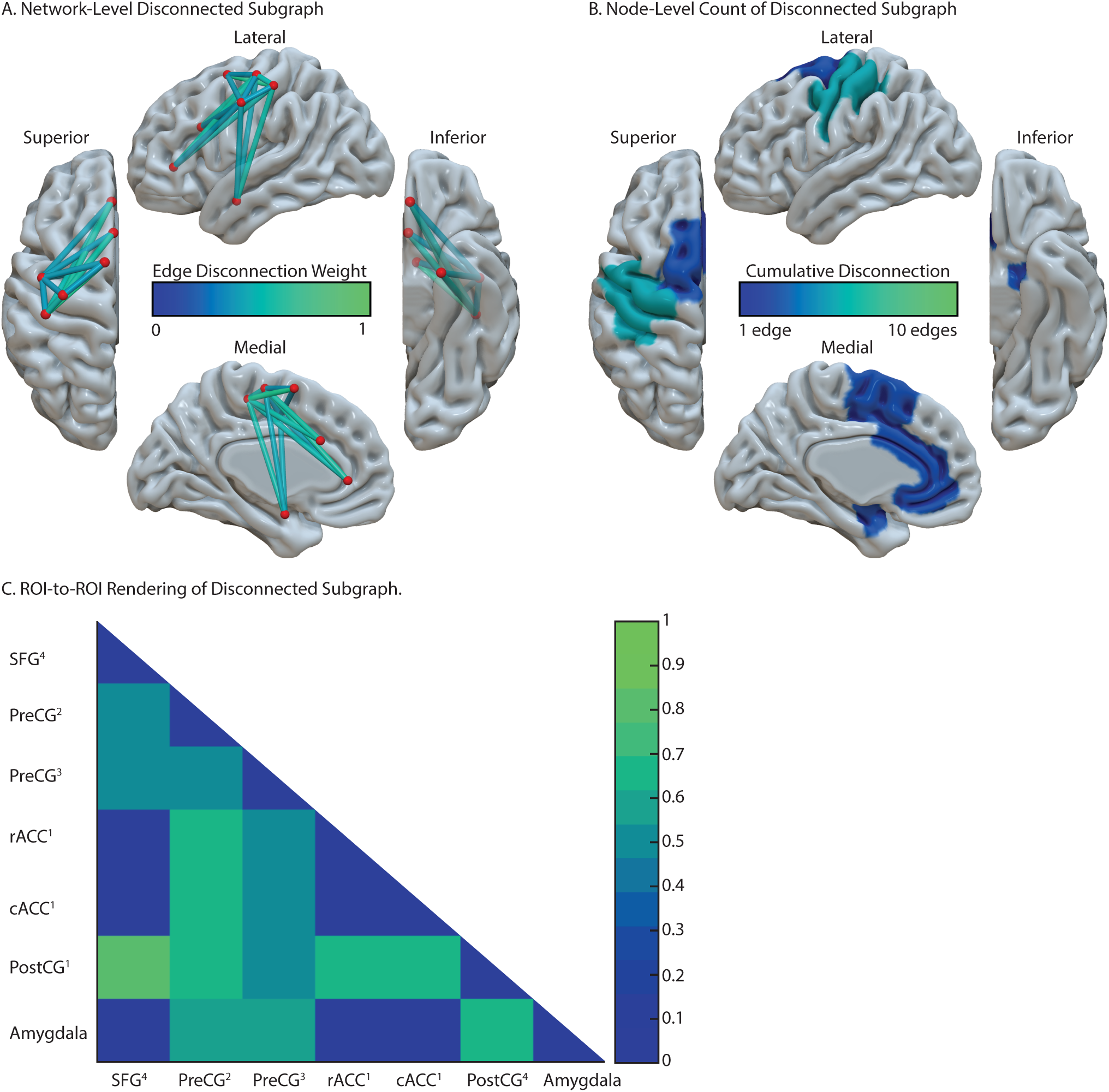
Support Vector Regression Connectome Lesion Symptom Mapping of Weakness in Right Hand Grip Strength. **A**. Maximally disconnected edges associated with reduced right-hand grip strength (weak grip strength with the right hand controlling for grip strength of the left hand and total lesion volume; blue-to-green scale). **B**. Cumulative node-level disconnection associated with reduced right-hand grip strength (blue-to-green scale) presented on the cortical surface. **C**. A heatmap of the disconnected subgraph depicts the ROI-to-ROI disconnection patterns projected in Fig. 4A. *Abbreviations*. SFG, superior frontal gyrus; PreCG, precentral gyrus; PostCG, postcentral gyrus; Rostral ACC, rostral anterior cingulate cortex; Caudal ACC, caudal anterior cingulate cortex. Numbers in superscript indicate anatomical subregions from the Lausanne atlas (for average MNI coordinates of each subregion, see Supplemental Table 1).

### Neuroimaging Acquisition

#### Acquisition of Anatomic Scans

MRI scans included whole-brain T1-weighted MR images collected on a 3T (Siemens Trio, Erlangen, Germany; repetition time = 1620 ms, echo time = 3.87 milliseconds, field of view = 192 × 256 mm, 1 × 1 × 1 mm voxels) or a 1.5T (Siemens Sonata, repetition time = 3,000 milliseconds, echo time = 3.54 milliseconds, field of view = 24 cm, 1.25 × 1.25 × 1.25 mm voxels) scanner, using an eight-channel or sixty-four channel head coil. Lesions were manually segmented on each LCVA participant’s high-resolution T1-weighted structural images. Lesioned voxels, consisting of both grey and white matter, were assigned a value of 1 and preserved voxels were assigned a value of 0. Binarized lesion masks were then registered to a standard template (Montreal Neurological Institute “Colin27”) using a symmetric diffeomorphic registration algorithm (Avants et al., 2008, www.picsl.upenn.edu/ANTS). Volumes were first registered to an intermediate template comprised of healthy brain images acquired on the same scanner. Volumes were then mapped onto the “Colin27” template to complete the transformation into standardized space. To ensure accuracy during the transformation process, lesion maps were subsequently inspected by a neurologist (H.B. Coslett), who was naïve to the behavioral data of the study. For increased accuracy, the pitch of the template was rotated to approximate the slice plane of each LCVA participant’s scan. This method has been demonstrated to achieve high intra- and inter-rater reliability (e.g., see Schnur et al., 2009). See Figure 1 for a rendering of each LCVA participant’s lesion and Table 1 for demographic information for each participant.

### Support Vector Regression Lesion Symptom Mapping Analyses

#### Support Vector Regression Voxel-based Lesion Symptom Mapping (SVR-VLSM)Analyses

SVR-VLSM was performed in MATLAB 2017B using a toolbox developed by DeMarco and Turkeltaub (2018) (https://github.com/atdemarco/svrlsmgui/). SVR-VLSM is a multivariate technique that uses machine learning to determine the association between lesioned voxels and behavior when considering the lesion status of all voxels submitted to the analysis. It overcomes several limitations of univariate VLSM, including inflated false positives from correlated neighboring voxels (Pustina et al., 2018), type II error due to correction for multiple comparisons (Bennett et al., 2009), and uneven statistical power due to biased lesion frequency as a function of vascular anatomy (Mah et al., 2014; Sperber & Karnath, 2017). SVR-VLSM has been shown to be superior to VLSM when multiple brain areas are involved in a single behavior (Herbet et al., 2015; Mah et al., 2014; Mirman, Zhang, et al., 2015; but see Sperber, 2019; Sperber et al., 2019; for discussion, see Zhang et al., 2014).

We performed two SVR-VLSM analyses. The first analysis tested the hypothesis that impaired tool use gesturing, controlling for variability in meaningless gesture imitation, would be associated with lesions to the left inferior parietal lobule, the left middle temporal gyrus, and left inferior and middle frontal gyri. The dependent measure was hand posture accuracy scores when gesturing tool use, residualized against hand posture accuracy scores when imitating meaningless gestures. A second SVR-VLSM analysis tested the hypothesis that reduced right-hand grip strength would be associated with lesions to the left primary motor cortex, premotor cortex, and subcortical structures including the basal ganglia. The dependent measure was right-hand grip strength score residualized against left-hand grip strength scores.

Only voxels lesioned in at least 10% of participants (5 participants in the analysis of tool use gesturing; 4 participants in the analysis of right-hand grip strength) were included. We controlled for variability in lesion volume using the ‘Direct Total Lesion Volume Control’ corrective method (see DeMarco & Turkeltaub, 2018). Five-fold cross-validation was implemented, in which 80% of the participants’ lesions and behavioral data were used to train a classifier, and the remaining 20% of participants’ lesions and behavioral data were used to test the classifier. This procedure was iterated 5 times to ensure that each unique subset of participant data was independently used for training and testing, and the resulting 5 maps of feature weights were averaged together to derive a final averaged map of voxelwise beta values. Voxelwise statistical significance was then determined using a Monte Carlo style permutation analysis in which the behavioral data were randomly assigned to a lesion map, and the same procedure as described above was iterated 10,000 times. Voxelwise z-scores were then computed for the true data in relation to the mean and standard deviation of voxelwise null distributions; the resulting z-score map was set to a threshold of z > 1.65 (*p* < .05, one-tailed) to determine chance-level likelihood of a lesion-symptom relation. We then further restricted the resulting map by eliminating any clusters with fewer than 500 contiguous voxels (see Garcea, Stoll, et al., 2019; Lacey et al., 2017; Skipper-Kallal et al., 2017). The Anatomical Automatic Labeling (AAL) atlas (Tzourio-Mazoyer et al., 2002) was used to assess overlap of significant voxels in the SVR-VLSM analyses with cortical and subcortical regions.

### Support Vector Regression Connectome-based Lesion Symptom Mapping

#### Support Vector Regression Connectome-based Lesion Symptom Mapping (SVR-CLSM)

We derived a whole-brain connectome—representing structural disconnection among cortical and subcortical nodes for each participant—to determine the group-level relation between node-to-node disconnection and impairment in tool use gesturing (e.g., see Gleichgerrcht et al., 2017).

Similar to SVR-VLSM, we used support vector regression in tandem with the disconnectome data to investigate connectome-wide disconnection in relation to cognitive impairment, which overcomes limitations of the univariate approach (e.g., see Pustina et al., 2017; Yourganov et al., 2016), and has implication for patient outcome prediction in acute stroke (Kuceyeski et al., 2016). First, each LCVA participant’s stroke lesion was drawn on their corresponding native T1-weighted MP-RAGE volume, and then normalized to a custom T1-weighted template constructed from 40 participants of the Human Connectome Project (HCP) (Greene et al., 2018) using cost function masking in ANTS (Avants et al., 2008). We then parcellated each participant’s MP-RAGE volume using the Lausanne 2008 atlas (Daducci et al., 2012; Hagmann et al., 2008). The Lausanne atlas contains 128 nodes distributed throughout the right and left hemispheres; for the purpose of the current investigation, we analyzed the 64 left-hemisphere nodes only, removing the left cerebellum (see Supplemental Table 1 for MNI coordinates of cortical and subcortical nodes). For any given node pair in the atlas, fractional connectivity loss was estimated by averaging the node-to-node connectivity loss due to each LCVA participant’s lesion relative to 210 neurotypical participants’ shortest path connectivity derived from the HCP dataset. The likelihood of node-to-node connectivity loss was then estimated on a participant-by-participant basis (Glasser et al., 2013). Specifically, the percent loss in structural connectivity between any two nodes was calculated as the fraction of the cumulative shortest path probability intersecting the lesion relative to the total shortest path probability shared between any given node pair in the HCP dataset (Greene et al., 2019), resulting in a participant-specific disconnectome.

We then conducted a group-level SVR-CLSM analysis in three steps. First, we transformed each participant’s whole-brain disconnectome into a vector, in which all right hemisphere nodes were zeroed out; we then combined each LCVA participant’s vector to form a group-level array. Next, edges were included in the analysis only if they were lesioned in at least 10% of participants, akin to the SVR-VLSM approach. Finally, in one statistical model we regressed variability in node-to-node disconnection on variability of three factors: **(1)** The number of months post-stroke of each LCVA participant; **(2)** Total lesion volume of each LCVA participant; and **(3)** Cumulative disconnection of the regions identified in the SVR-VLSM analysis (see Table 2).

**Table 2.** Peak MNI coordinates identified in the SVR-VLSM analysis of errors in tool use gesturing controlling for errors in meaningless imitation (A), and reduced right-hand grip strength controlling for a reduction in left-hand grip strength (B). Region labels were derived from the Automated Anatomical Labeling (AAL) template.

| **A. Errors in Tool Use Gesturing, Controlling for Errors in Meaningless Imitation** |  |  |  |  |  |  |
| --- | --- | --- | --- | --- | --- | --- |
| AAL Label | AAL Number | Mean Center of Mass (XYZ) |  |  | Peak Z-Score | Number of Voxels |
| Left Precentral Gyrus | 1 | -36 | 11 | 39 | -3.72 | 4937 |
| Left Superior Frontal Gyrus | 3 | -20 | 22 | 44 | -2.48 | 30 |
| Left Middle Frontal Gyrus | 7 | -39 | 13 | 38 | -3.72 | 5593 |
| Left Pars Opercularis | 11 | -52 | 22 | 34 | -3.55 | 3475 |
| Left Pars Triangularis | 13 | -56 | 18 | 30 | -3.13 | 4284 |
| Left Rolandic Operculum | 17 | -52 | 6 | 9 | -2.95 | 289 |
| Left Insula | 29 | -30 | 23 | 16 | -2.04 | 59 |
| Left Superior Occipital Gyrus | 49 | -22 | -73 | 25 | -2.84 | 91 |
| Left Middle Occipital Gyrus | 51 | -35 | -65 | 27 | -3.36 | 962 |
| Left Postcentral Gyrus | 57 | -34 | -27 | 41 | -3.02 | 3316 |
| Left Superior Parietal Lobule | 59 | -27 | -47 | 51 | -2.45 | 162 |
| Left Inferior Parietal Lobule | 61 | -52 | -30 | 41 | -3.72 | 4621 |
| Left Supramarginal Gyrus | 63 | -63 | -43 | 25 | -3.13 | 1232 |
| Left Angular Gyrus | 65 | -42 | -58 | 24 | -3.44 | 2244 |
| Left Caudate Nucleus | 71 | -19 | 19 | 17 | -1.89 | 6 |
| Left Heschel's Gyrus | 79 | -36 | -29 | 9 | -3.07 | 175 |
| Left Superior Temporal Gyrus | 81 | -44 | -26 | 5 | -3.72 | 2288 |
| Left Middle Temporal Gyrus | 85 | -46 | -58 | 22 | -3.55 | 717 |
| **B. Weakened Right-Hand Grip Strength, Controlling for Left-Hand Grip Strength** |  |  |  |  |  |  |
| AAL Label | AAL Number | Mean Center of Mass (XYZ) |  |  | Peak Z-Score | Number of Voxels |
| Left Precentral Gyrus | 1 | -41 | -3 | 27 | -2.49 | 720 |
| Left Middle Frontal Gyrus | 7 | -32 | 14 | 37 | -2.22 | 17 |
| Left Rolandic Operculum | 17 | -51 | -3 | 16 | -2.49 | 264 |
| Left Insula | 29 | -34 | -2 | 15 | -2.79 | 1586 |
| Left Amygdala | 41 | -29 | -6 | -10 | -1.77 | 3 |
| Left Postcentral Gyrus | 57 | -39 | -17 | 42 | -2.55 | 2213 |
| Left Superior Parietal Lobule | 59 | -35 | -40 | 57 | -1.66 | 2 |
| Left Inferior Parietal Lobule | 61 | -45 | -25 | 41 | -1.68 | 20 |
| Left Caudate Nucleus | 71 | -15 | 6 | 18 | -2.62 | 194 |
| Left Putamen | 73 | -29 | -2 | 13 | -2.75 | 2095 |
| Left Pallidum | 75 | -21 | 2 | 7 | -1.99 | 29 |
| Left Superior Temporal Gyrus | 81 | -43 | -10 | 1 | -1.95 | 1 |

In the analysis of tool use gesturing, cumulative lobe-level structural disconnection was calculated for the left frontal cortex (inferior, middle, superior frontal gyri), the left temporal lobe (superior temporal gyrus, middle temporal gyrus, transverse temporal, bank of the superior temporal sulcus), and the left parietal lobe (supramarginal gyrus, angular gyrus, superior parietal lobule). Variability in lobe-level disconnection was regressed from variability in node-to-node disconnection. Similarly, in the analysis of right-hand grip strength, cumulative structural disconnection was calculated for the left cortical (pre- and post-central gyri) and subcortical motor system (left insula, and left subcortical regions including the thalamus, caudate, putamen, pallidum, and nucleus accumbens), and variability in cumulative disconnection was regressed from variability in node-to-node disconnection. In this way, we use the SVR-VLSM results as a ‘localizer’ map to independently identify regions of relevance in the SVR-CLSM analysis, and by regressing cumulative structural disconnection of those sites on node-to-node disconnection variability, we ensure that our observed disconnection patterns exist over and above cumulative disconnection of a given node contributing to the behavioral task of relevance. Given that there was a moderate relation between months post-onset and lesion volume in the full group of LCVA participants (r(64) = 0.22, *p* = 0.07), we regressed variability in lesion volume from variability in months post-onset, and used residual months post-onset values in the model described above.

The residual structural connectivity values were then entered in a support vector regression algorithm in MATLAB 2017B (function: ‘fitrsvm’) using a Gaussian kernel function (‘rbf’) and the default ‘auto’ kernel scale. Cross-validation was used, in which 80% of the disconnection data were used to train the model to learn the relation between disconnection and behavioral scores, and 20% of the disconnection data were used to test the robustness of that model using a left-out sample. After iterating this procedure 5 times, all data were used to independently train and test the classifier, which resulted in 5 disconnectome maps of feature weights (beta values), indicating the strength of a given edge predicting behavioral scores. We then averaged across the 5 disconnectome maps to obtain a final map, and transformed the map into a 2-dimensional representation for visualization (64×64 matrix of left hemisphere disconnection).

To interpret the strength of edge-level beta weights, we conducted a Monte Carlo style permutation analysis in which we randomly assigned behavioral scores to disconnectome maps, and repeated the analysis 10,000 times; all other aspects of the permutation analysis were identical to the analysis of the true data. The result of the permutation analysis is a distribution of beta weights for a given edge that arise due to chance; we then z-score the beta weights of the true data relative to the mean and standard deviation of the null distribution. The resulting z-value matrix was set to a threshold of z = 1.65 (*p* < .05, one-tailed) to identify above-chance beta values.

#### Calculating Maximally Disconnected Subgraphs

Because the full disconnectome can be quite extensive with many region pairs having negligible connectivity loss (rendering visualization and interpretation challenging), we reduce the dimensionality of the z-score disconnectome by extracting a maximally disconnected subgraph. The subgraph is extracted by initializing a subgraph with the brain regions that share the edges with the greatest connectivity loss; critically, only edges that survive Monte Carlo permutation analysis (z < −1.65, one-tailed) are entered into the subgraph analysis; all non-significant values are zeroed-out, ensuring that the subgraph analysis identifies regions for which disconnection is significantly associated with worse performance on the test of interest. Subsequently, regions are added such that the cumulative amount or weight of the connectivity loss is maximized amongst the regions; this procedure identifies the maximally disconnected subgraph when the change in the weight contributed by each additional region added peaks. The maximally disconnected subgraph contains regions and edges with the greatest shared connectivity loss associated with reduced performance on the test of interest (for precedent in the analysis of stroke disconnection, see Greene et al., 2019).

## Results

### Support Vector Regression Voxel-based Lesion Symptom Mapping Results

Figure 2A depicts lesion overlap among the 57 participants with high resolution MRI anatomical data (see Supplemental Figure 1 for a rendering of structural disconnection overlap among the 57 participants). Two SVR-VLSM analyses were conducted; in the first analysis we identified voxels in which lesions were associated with worse performance when gesturing tool use to the sight of objects (controlling for variability in meaningless imitation). In the second analysis we identified voxels in which lesions were associated with reduced right-hand grip strength (controlling for variability in left-hand grip strength).

#### Tool Use Gesturing

Lesions to two large clusters were associated with poor tool use gesturing performance (Figure 2B). The first cluster included the left inferior parietal lobule (left supramarginal gyrus, left angular gyrus), the left superior parietal lobule, the left superior temporal gyrus, the left middle temporal gyrus, and posterior voxels including portions of middle and superior occipital gyri (see Table 2A). A second cluster identified voxels in left frontal cortex, including pars opercularis, pars triangularis, the middle frontal gyrus, and the superior frontal gyrus. This cluster extended posteriorly to include the left pre-central gyrus, the left insula, and the left caudate nucleus (see Table 2A). Our findings replicate prior univariate work (Buxbaum et al., 2014; Watson & Buxbaum, 2015) using a multivariate approach (see Supplemental Figure 2).

#### Reduced Grip Strength

Lesions to two clusters were associated with reduced right-hand grip strength (see Figure 2C). The first cluster included pre- and post-central gyri, the left inferior and superior parietal lobule, and the left middle frontal gyrus. A second cluster included medial portions of the left superior temporal gyrus, the left insula, basal ganglia structures (including the putamen, caudate nucleus, and pallidum), and the left amygdala (see Table 2B). These results confirm that reduced right-hand grip strength is associated with lesions to the cortical motor system and to additional regions including basal ganglia and insula, largely bypassing the lesion sites associated with reduced tool use gesturing.

### Support Vector Regression Connectome-based Lesion Symptom Mapping Results

#### Tool Use Gesturing Maximally Disconnected Subgraph

As shown in Figure 3A, there were three clusters identified as maximally disconnected in association with reduced tool use gesturing (controlling for variability in meaningless imitation and total lesion volume). First, we identified inferior-going disconnection between the left inferior parietal lobule and superior temporal sulcus, middle temporal gyrus, lingual gyrus, parahippocampal gyrus, and hippocampus. Second, we observed disconnection among pre- and post-central gyri, middle frontal gyrus, superior frontal gyrus, and the inferior parietal lobule. Third, the superior temporal gyrus was associated with disconnection to all nodes except pre-and post-central gyri (see Figure 3A, and Table 3). Aggregating node-level disconnection resulted in a map of cumulative disconnection loss, which identifies the superior temporal sulcus and the inferior parietal lobule as the most disconnected nodes associated with low tool use gesturing scores (see Figure 3B and Table 3). Node-to-node disconnection values are reported in Figure 3C. For a rendering of the full, uncorrected z-score matrix, see Supplemental Figure 3.

**Table 3.** MD Subgraph from SVR-CLSM of hand posture errors when gesturing tool use. Feature weights range from −0.32 to −0.72; all edges survive Monte Carlo style permutation analysis (10,000 iterations) with z-scores less than or equal to −1.65 (*p* < .05, one-tailed). *Abbreviations*. rMFG, rostral middle frontal gyrus; SFG, superior frontal gyrus; PreCG, precentral gyrus; PostCG, postcentral gyrus; IPL, inferior parietal lobule; LG, lingual gyrus; PPG, parahippocampal gyrus; MTG, middle temporal gyrus; B. STS, bank of the superior temporal sulcus; STG, superior temporal gyrus; TT, transverse temporal. Numbers in subscript indicate anatomical subregions from the Lausanne atlas (for the average MNI coordinates of each subregion, see Supplemental Table 1).

|  | rMFG<sup>1</sup> | rMFG<sup>2</sup> | SFG<sup>1</sup> | SFG<sup>2</sup> | Pre<br>CG<sup>4</sup> | Post<br>CG<sup>1</sup> | IPL<sup>2</sup> | LG<sup>1</sup> | LG<sup>2</sup> | PPG<sup>1</sup> | MTG<sup>1</sup> | B.STS<sup>1</sup> | STG<sup>1</sup> | TT1 | Hippocampus |
| --- | --- | --- | --- | --- | --- | --- | --- | --- | --- | --- | --- | --- | --- | --- | --- |
| rMFG<sup>1</sup> | — |  |  |  |  |  |  |  |  |  |  |  |  |  |  |
| rMFG<sup>2</sup> | — | — |  |  |  |  |  |  |  |  |  |  |  |  |  |
| SFG<sup>1</sup> | — | — | — |  |  |  |  |  |  |  |  |  |  |  |  |
| SFG<sup>2</sup> | — | — | — | — |  |  |  |  |  |  |  |  |  |  |  |
| PreCG<sup>4</sup> | -0.48 | -0.41 | -0.47 | -0.42 | — |  |  |  |  |  |  |  |  |  |  |
| PostCG<sup>1</sup> | -0.55 | -0.54 | -0.55 | -0.64 | — | — |  |  |  |  |  |  |  |  |  |
| IPL<sup>2</sup> | — | — | -0.36 | -0.33 | — | — | — |  |  |  |  |  |  |  |  |
| LG<sup>1</sup> | — | — | — | — | — | — | — | — |  |  |  |  |  |  |  |
| LG<sup>2</sup> | — | — | — | — | — | — | -0.32 | — | — |  |  |  |  |  |  |
| PPG<sup>1</sup> | — | — | — | — | — | — | -0.39 | — | — | — |  |  |  |  |  |
| MTG<sup>1</sup> | — | — | — | — | — | — | -0.39 | — | — | — | — |  |  |  |  |
| B.STS<sup>1</sup> | -0.40 | -0.42 | -0.45 | -0.47 | — | — | -0.41 | -0.68 | -0.72 | -0.61 | -0.56 | — |  |  |  |
| STG<sup>1</sup> | -0.53 | -0.40 | -0.39 | -0.48 | — | — | — | -0.60 | -0.57 | -0.43 | -0.59 | -0.67 | — |  |  |
| TT<sup>1</sup> | -0.52 | — | — | — | — | — | — | — | — | — | -0.47 | -0.60 | -0.50 | — |  |
| Hippocampus | — | — | — | — | — | — | -0.27 | — | — | — | — | -0.57 | -0.37 | — | — |
|  | rMFG<sup>1</sup> | rMFG<sup>2</sup> | SFG<sup>1</sup> | SFG<sup>2</sup> | Pre<br>CG<sup>4</sup> | Post<br>CG<sup>1</sup> | IPL<sup>2</sup> | LG<sup>1</sup> | LG<sup>2</sup> | PPG<sup>1</sup> | MTG<sup>1</sup> | B.STS<sup>1</sup> | STG<sup>1</sup> | TT1 | Hippocampus |
| Cumulative<br>Disconnection | 5 | 4 | 5 | 5 | 4 | 4 | 7 | 2 | 3 | 3 | 4 | 12 | 11 | 4 | 3 |

#### Reduced RH Grip Strength Maximally Disconnected Subgraph

As shown in Figure 4A, pre- and post-central gyri were identified as maximally disconnected in association with reduced right-hand grip strength (controlling for left-hand grip strength and total lesion volume; see Table 4). This analysis identified the superior frontal gyrus, rostral and caudal anterior cingulate cortices, and the amygdala as nodes disconnected from pre- and post-central gyri (see Figure 4B for node-level count, and Figure 4C for a visualization of node-to-node disconnection). For a rendering of the full, uncorrected z-score matrix, see Supplemental Figure 4.

**Table 4.** MD Subgraph from SVR-CLSM of weak right-hand grip strength controlling for left-hand grip strength. Feature weights range from −0.50 to −0.81; all edges survive Monte Carlo style permutation analysis (10,000 iterations) with z-scores less than or equal to −1.65 (*p* < .05, one-tailed). *Abbreviations*. SFG, superior frontal gyrus; PreCG, precentral gyrus; PostCG, postcentral gyrus; Rostral ACC, rostral anterior cingulate cortex; Caudal ACC, caudal anterior cingulate cortex. Numbers in superscript indicate anatomical subregions from the Lausanne atlas (for average MNI coordinates of each subregion, see Supplemental Table 1).

|  | SFG<sup>4</sup> | Pre CG<sup>2</sup> | Post CG<sup>3</sup> | Rostral ACC<sup>1</sup> | Caudal ACC<sup>1</sup> | Post CG<sup>1</sup> | Amygdala |
| --- | --- | --- | --- | --- | --- | --- | --- |
| SFG<sup>4</sup> | — |  |  |  |  |  |  |
| Pre CG<sup>2</sup> | -0.52 | — |  |  |  |  |  |
| Post CG<sup>3</sup> | -0.54 | -0.51 | — |  |  |  |  |
| Rostral ACC<sup>1</sup> | — | -0.70 | -0.54 | — |  |  |  |
| Caudal ACC<sup>1</sup> | — | -0.69 | -0.53 | — | — |  |  |
| Post CG<sup>1</sup> | -0.81 | -0.64 | -0.52 | -0.63 | -0.67 | — |  |
| Amygdala | — | -0.56 | -0.57 | — | — | -0.66 | — |
|  | SFG<sup>4</sup> | Pre CG<sup>2</sup> | Post CG<sup>3</sup> | Rostral ACC<sup>1</sup> | Caudal ACC<sup>1</sup> | Post CG<sup>1</sup> | Amygdala |
| Cumulative Disconnection | 3 | 6 | 6 | 3 | 3 | 6 | 3 |

### Post-hoc Analysis of Left Inferior Parietal Lobule Disconnection

Prior work has identified reduced functional connectivity between the left inferior parietal lobule and left medial fusiform gyrus in association with the severity of tool use gesturing, yet we did not observe this pattern in our analysis of maximal disconnection associated with tool use gesturing scores (Figure 3). We therefore conducted a post-hoc analysis using linear regression to investigate left inferior parietal disconnectivity. The procedure was carried out in three steps. First, as described above, we used residual disconnection values after removing variability in disconnection associated with months post-onset, total lesion volume, and cumulative disconnection of nodes identified in the SVR-VLSM analysis. We then computed the correlation between disconnection of the left inferior parietal lobule (identified in the maximally disconnected subgraph analysis) with all other left hemisphere nodes, and imposed three criteria to determine the significance of disconnection in association with severity of tool use gesturing performance.

First, we inspected connectivity patterns only if the node under consideration was outside of the lesion territory (see Figure 2A for lesion overlap map). We did this by identifying candidate nodes only if there was less than 1% voxelwise overlap with the voxels in the remote node and the lesion overlap map. This first criterion ensures that parietal disconnection to remote nodes cannot be due to weak signal at the remote site. Second, the connectivity relation (correlation value) with the left inferior parietal node was considered only if it was significant (minimum r < - 0.26, *p* > .05). Third, we then conducted a Monte Carlo style permutation analysis in which we randomly assigned the behavioral data to the disconnection data using 10,000 iterations to derive a null distribution from which to z-score the true data. Thus, all disconnection to the inferior parietal node **(1)** Needed to exist outside of the lesion territory; **(2)** Needed to predict a significant amount of variance in tool use gesturing performance (controlling for meaningless gesture imitation performance and total lesion volume); and **(3)** Needed to be at least 2 standard deviations (Z < - 1.96) outside the mean of a null distribution for a given edge as determined by Monte Carlo style permutation analysis.

Figure 5 depicts the result of this analysis. We found significant and robust disconnection between the left inferior parietal lobule and **(i)** Ventral temporal regions including fusiform gyrus, lingual gyrus, parahippocampal gyrus, and calcarine cortex extending medially into the cuneus; and **(ii)** Between the inferior parietal lobule and frontal and medial regions including the superior frontal gyrus and anterior cingulate, respectively (see Figure 5A; see Table 5 for correlation values and z-score values). The node-level rendering of inferior parietal disconnection depicts the full extent of ventral temporal cortex and medial-frontal involvement (Figure 5B).

**Table 5.** Analysis of inferior parietal lobule disconnection in relation to hand posture errors in tool use gesturing, controlling for hand posture errors in meaningless imitation and total lesion volume. Numbers in superscript indicate anatomical subregions from the Lausanne atlas (for average MNI coordinates of each subregion, see Supplemental Table 1).

|  | Correlation Value | Significance Value | Z-score |
| --- | --- | --- | --- |
| Superior Frontal Gyrus <sup>1</sup> | -0.33 | 0.01 | -2.39 |
| Superior Frontal Gyrus <sup>2</sup> | -0.28 | 0.03 | -2.12 |
| Isthmus Cingulate <sup>1</sup> | -0.29 | 0.03 | -2.18 |
| Cuneus <sup>1</sup> | -0.27 | 0.04 | -2.03 |
| Pericalcarine <sup>1</sup> | -0.27 | 0.04 | -2.06 |
| Lingual Gyrus <sup>1</sup> | -0.28 | 0.03 | -2.14 |
| Lingual Gyrus <sup>2</sup> | -0.28 | 0.03 | -2.13 |
| Fusiform Gyrus <sup>1</sup> | -0.27 | 0.05 | -1.99 |
| Parahippocampal Gyrus <sup>1</sup> | -0.34 | 0.01 | -2.54 |

**Figure 5.**
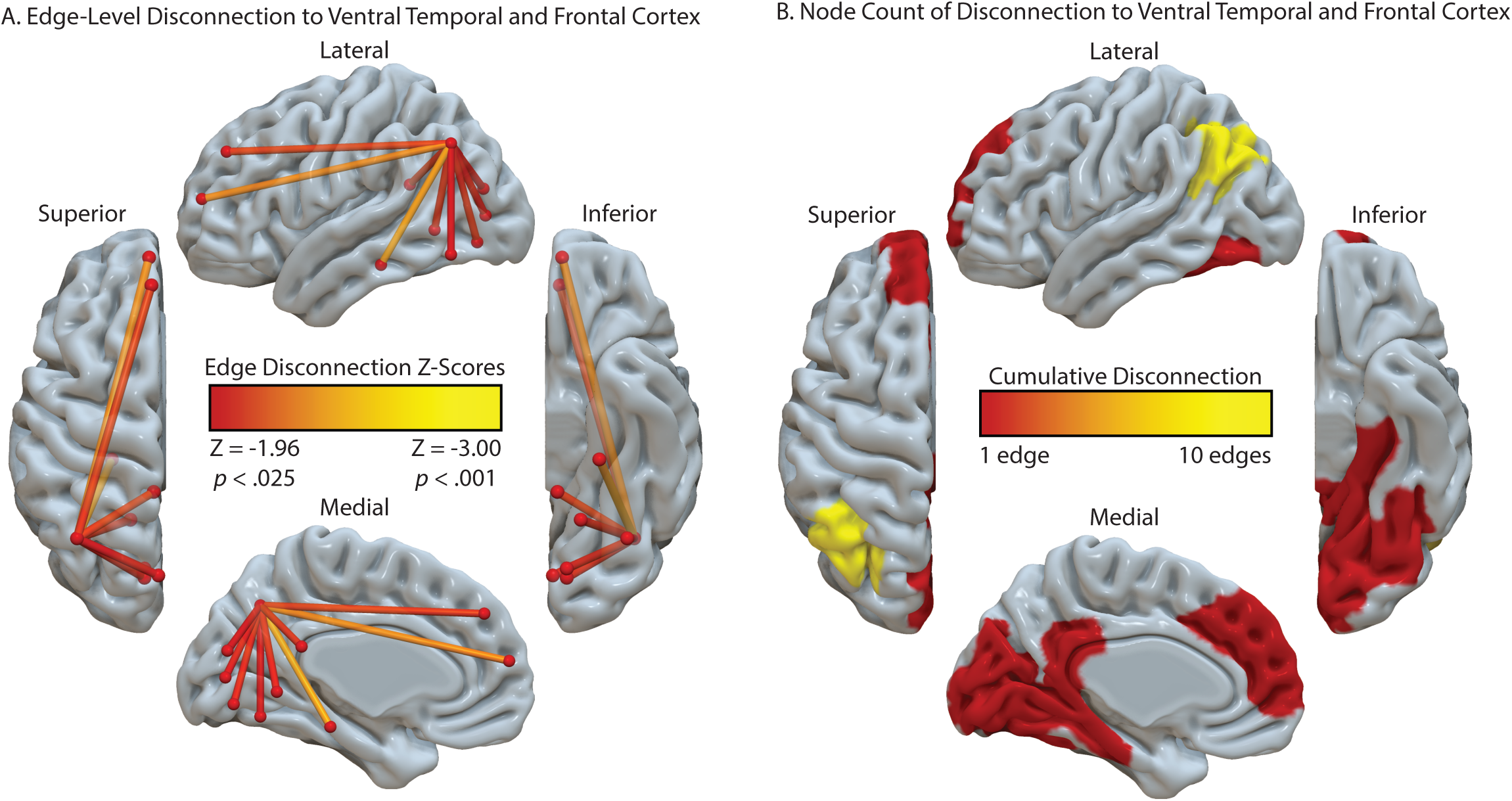
Analysis of IPL Disconnection Associated with Errors in Tool Use Gesturing. **A**. Edge-level disconnection identifies the left ventral temporal cortex and medial frontal regions as sites that exhibit a disconnection to the left inferior parietal lobule. Regions are included in the analysis if **(1)** The region is outside of the lesion territory (< 1% overlap with the lesion overlap map (see Figure 2A)), **(2)** The correlation between disconnection and tool use gesturing is significant, and **(3)** The magnitude of the correlation is at least 2 standard deviations above the mean of a null distribution derived from a Monte Carlo style permutation analysis (10,000 iterations). **B**. Cumulative node-level disconnection associated with reduced tool use gesturing performance is projected on the cortical surface.

## General Discussion

Prior work has identified the Tool Use Network—a whole-brain network of regions working in concert to support the retrieval of action and object knowledge in the service of implementing object-directed action. We performed SVR-VLSM and SVR-CLSM analyses with 57 LCVA participants to elucidate the relation among tool use gesturing performance, lesion location, and structural disconnection. To our knowledge, this is the first study to combine lesion- and connectome-based symptom mapping in tandem with support vector machine learning to investigate local and non-local effects of LCVA lesions on tool use ability in apraxia. Moreover, the analysis of reduced contralesional grip strength permitted us to test the specificity of disconnection within the Tool Use Network as a key substrate that gives rise to reduced tool use gesturing, and not reduced grip strength.

Consistent with the predictions of neurocognitive models of praxis (e.g., see Binkofski & Buxbaum, 2013; Buxbaum, 2017), impairments in tool use gesturing were associated with lesions in the left inferior parietal lobule, the left inferior and middle frontal gyri, and the left posterior middle temporal gyrus that extended into ventral occipitotemporal cortex. These results are in agreement with prior univariate VLSM analyses, including results from our lab (Buxbaum et al., 2014; Tarhan et al., 2015; Watson & Buxbaum, 2015), and other key findings in the literature (Dressing et al., 2018; Haaland et al., 2000; Manuel et al., 2013; M. Martin et al., 2016). In contrast, a reduction in right-hand grip strength, controlling for shared variance in left-hand grip strength, was associated with lesions in the left pre- and post-central gyri, the left insula, and in voxels extending subcortically to include regions of the basal ganglia (putamen, pallidum, caudate nucleus). These findings are in accord with prior univariate VLSM analyses of reduced contralesional grip strength (Goldenberg & Spatt, 2009; Greene et al., 2019), and demonstrate that the lesion sites associated with reduced strength of the right hand minimally overlap with nodes of the Tool Use Network.

We reasoned that tool use gesturing ability would be associated with the degree of disconnection among nodes of the Tool Use Network, controlling for shared variance when imitating meaningless gestures. Our SVR-CLSM analysis revealed that the degree of hand posture errors when gesturing tool use was associated with disconnection among nodes in the left inferior parietal lobule, the left superior frontal gyrus, the left middle temporal gyrus, and left ventral (lingual gyrus, fusiform gyrus) and mesial (parahippocampal gyrus, hippocampus) temporal cortices. Finally, voxels in the temporo-parietal junction (superior temporal sulcus, inferior parietal lobule) were strongly disconnected from nearly all regions identified, indicating that damage to the white matter adjacent to temporo-parietal cortex has long-range disconnective effects. This is due, in part, to the proximity of white matter pathways medial to the superior temporal gyrus, an issue we return to below.

We then used SVR-CLSM over right-hand grip strength data to test the hypothesis that reductions in contralesional grip strength would be associated with disconnection among nodes of the cortical and subcortical motor system, bypassing nodes of the Tool Use Network. Consistent with our hypothesis, we found reduced right-hand grip strength in association with disconnection among pre- and post-central gyri, and superior frontal gyrus, anterior cingulate cortex, and the amygdala. Critically, there was minimal overlap between nodes of the Tool Use Network and regional disconnection associated with reduced right-hand grip strength.

### Implications for Neurocognitive Models of Praxis in the Human Brain

We observed that structural disconnection with the left inferior parietal lobule was associated with reduced tool use gesturing performance in apraxia. The left inferior parietal lobule forms a core component of the ventro-dorsal stream, a visuomotor processing pathway that supports the retrieval of manipulation knowledge for tool use, interfacing current visual input with conceptual representations of actions (Binkofski & Buxbaum, 2013). In neurotypical adults, diffusion tractography studies have identified the requisite connectivity to interface the left inferior parietal lobule with other nodes in the Tool Processing Network. For example, posterior fibers of the arcuate fasciculus (pAF) provide the anatomical substrate to connect the lateral and posterior middle temporal gyrus with the left inferior parietal lobule (Ramayya et al., 2010), and structural connectivity between the left inferior parietal lobule and frontal-motor sites, subserved by fibers of the superior longitudinal fasciculus (SLF), is critical for action selection and planning (Caspers et al., 2011; Ramayya et al., 2010; Ruschel et al., 2014; Rushworth et al., 2006). Furthermore, lesions involving the SLF are associated with poor tool use gesturing in apraxia (Bi et al., 2015; Watson & Buxbaum, 2015), as well as impairments in word repetition (Chernoff et al., 2020), and increased phonological errors in picture naming (Schwartz et al., 2012), suggesting the SLF underlies sensory-to-motor mappings across language and action domains.

In contrast, the ventral fiber pathway runs inferior to the superior temporal gyrus, coursing anteriorly and ventrally through fibers of the extreme capsule and uncinate fasciculus, and is argued to play a domain-general role in conceptual processing across numerous cognitive domains, including tool use pantomiming (Vry et al., 2015) and language (Saur et al., 2008; Weiller et al., 2011). Our analyses did not identify regions connected by the ventral pathway (e.g., the left inferior frontal gyrus), suggesting that the integrity of this tract is not causally related to tool use gesturing in apraxia. Given prior evidence that lesions involving the white matter adjacent to the inferior frontal gyrus (uncinate fasciculus, left inferior fronto-occipital fasciculus, anterior thalamic radiations) were associated with impaired selection of verbal and non-verbal conceptual knowledge (e.g., see Mirman, Chen, et al., 2015; see also Han et al., 2013), it remains a possibility that the ventral fiber pathway supports tool action selection. However, given that current lesion evidence suggests that action selection in tool use is mediated by fronto-parietal structures via the SLF (e.g., see Garcea, Stoll, et al., 2019; Watson & Buxbaum, 2015), it will be important for future VLSM and CLSM work to determine the extent to which damage to the ventral fiber pathway, controlling for damage of SLF fibers, is associated with action selection difficulties.

More recently, it has been argued that fibers of the vertical occipital fasciculus (VOF) provide a substrate to connect the ventral and dorsal object processing pathways (Yeatman et al., 2014). The VOF runs lateral to the inferior longitudinal fasciculus and posterior to the arcuate fasciculus, connecting ventral occipitotemporal cortex with the posterior and inferior parietal lobule (Weiner et al., 2017). Though the integrity of VOF fibers is implicated in visual processing of faces (Weiner et al., 2016), written text (Yeatman et al., 2013), and objects (Freud et al., 2016), Budisavljevic and colleagues (2018) recently demonstrated that the speed with which participants reached maximum grip aperture when grasping an object was predicted by structural integrity of the VOF. Specifically, participants’ faster opening of the hand when grasping an object was associated with faster transfer of information between visual perceptual processing (analysis of shape, form, and surface texture; ventral stream) and object-directed grasping (dorsal stream). Our post-hoc analysis identified structural disconnection between the left inferior parietal lobule and ventral occipitotemporal cortex, including the left medial fusiform gyrus, in association with reduced tool use gesturing. It therefore remains a possibility that lesions to the posterior inferior parietal lobule damages superior terminations of the VOF, which subsequently gives rise to tool use gesturing impairment in apraxia. Although this speculation needs to be evaluated in future empirical work, prior studies in neurotypical participants have demonstrated increased functional connectivity between the inferior parietal lobule and left medial fusiform gyrus in tool and action processing (Assmus et al., 2007; Garcea et al., 2018; Garcea & Mahon, 2014), and recent lesion evidence suggests that inferior parietal-to-medial fusiform connectivity disruption predicts abnormal tool processing (Garcea, Almeida, et al., 2019) and tool use gesturing ability (Watson et al., 2019).

To bring our results into register with the apraxia literature, tool use gesturing performance is associated with distributed structural connectivity among the left inferior parietal lobule and lateral and inferior temporal cortices. Voxels in lateral occipitotemporal cortex respond to images of hands (Orlov et al., 2010) and tools (Bracci et al., 2012), and exhibit functional connectivity to the left inferior parietal lobule (Bracci et al., 2012; Mahon et al., 2007). Furthermore, parietal disconnection extended to ventral temporal cortex to include the fusiform gyrus, consistent with prior functional connectivity findings in apraxia (Watson et al., 2019) and in neurotypical adults (Garcea & Mahon, 2014; see also Chen et al., 2017; Stevens et al., 2015). In light of these findings, it has been argued that the processing of object properties in ventral temporal cortex provides an input to the praxis system, because the extraction of visual attributes of objects (e.g., whether an object is slippery, hot, or sharp) as well as stored knowledge of material properties (e.g., object weight; Gallivan et al., 2014) informs the retrieval of hand posture and deployment of an object-directed grasp when functionally manipulating a tool (Almeida et al., 2013; Mahon et al., 2013; for discussion, see Gallivan & Culham, 2015).

These action and object retrieval processes will, by hypothesis, interface with frontal-motor structures of relevance for action selection and motor implementation. We found that structural connectivity among the left inferior parietal lobule, left superior temporal gyrus, and middle and superior frontal gyri was strongly predictive of tool use gesturing ability. Prior fMRI work has implicated dorso-lateral prefrontal regions in attentional control and response selection (Cieslik et al., 2015), and lesions to middle and inferior frontal gyri are associated with tool use pantomiming deficits (Goldenberg et al., 2007; Haaland et al., 2000; Watson & Buxbaum, 2015; see also Bohlhalter et al., 2011). Considering that numerous fMRI studies have reported increased BOLD contrast in the inferior frontal gyrus when gesturing tool use (e.g., see Vry et al., 2015), it is surprising that the maximally disconnected subgraph analysis did not reveal increased disconnection with the left inferior frontal gyrus^1^. Although there is a paucity of structural connectivity research in apraxia, our results are consistent with Bi and colleagues’ (2015) finding that the severity of tool use ability was associated with structural disconnection between dorsolateral frontal cortex and pre-motor cortex. Their dorsolateral frontal area was in close anatomical proximity to the superior frontal gyrus that we identified as disconnected with the inferior parietal lobule (see Figure 3), suggesting that fronto-parietal damage has detrimental effects upon action selection and gesture implementation.

### Limitations

Whereas our SVR-VLSM result is consistent with a large body of work demonstrating that apraxia is associated with lesions to the supramarginal gyrus and angular gyrus (for review, see Buxbaum & Randerath, 2018; Goldenberg, 2009; Johnson-Frey, 2004), our SVR-CLSM analysis identified the posterior inferior parietal lobule in the vicinity of the left angular gyrus. This may be driven in part by the use of shortest path tractography, which identifies the optimal path between nodes such that the probability that adjacent voxels form a contiguous, short path between nodes is high. Visual inspection of the left inferior parietal lobule sub-region confirms its anatomical proximity to posterior fibers of the superior longitudinal fasciculus (see Supplemental Figure 5), indicating that lesions invading white matter adjacent to the posterior inferior parietal lobule are likely to disconnect the inferior parietal lobule from ventral and lateral temporal cortices. Importantly, prior work indicates that fibers of the VOF lie posterior to fibers of the AF/SLF (Weiner et al., 2017), which is also anatomically proximal to white matter voxels adjacent to the posterior inferior parietal lobule area identified in our analysis (e.g., see Jitsuishi et al., 2020). Thus, it is not surprising the posterior portion of the left inferior parietal lobule was identified as disconnected from lateral and ventral temporal cortices in association with tool use gesturing performance given its location to these white matter tracts.

A strength of our SVR-CLSM approach is that we control for the cumulative amount of node-level disconnection using the results of the SVR-VLSM. Thus, our SVR-CLSM findings are significant over and above total lesion volume, cumulative node-level disconnection, and months post-onset, and converge with the lesion sites identified in our SVR-VLSM analysis, giving us confidence that we are identifying a robust effect. Moving forward, it will be critical to use similar approaches to address recent critiques of multivariate lesion-symptom mapping, including issues of inadequate statistical control of non-uniform LCVA lesion distributions, and disagreement over the proper multiple comparison correction (for discussion, see DeMarco & Turkeltaub, 2018; Sperber & Karnath, 2018; Sperber et al., 2019). In the degree to which convergence across independent analyses can be demonstrated, we can start to develop analytic pipelines to control for lesions distributions and multiple comparison correction. In particular, graph theoretic algorithms like the maximally disconnected subgraph analysis we have used offer a data-driven approach to identify nodes that contribute most heavily to cumulative disconnection (see Greene et al., 2019).

Finally, in this study we focused solely on patterns of disconnection within the left hemisphere. It will be important for future studies to determine the extent to which apraxia severity is associated with interhemispheric disconnection following LCVA. Prior resting state functional connectivity studies demonstrate that recovery of motor and attentional processes is dependent on the integrity of interhemispheric functional connectivity (Carter et al., 2010; He et al., 2007; Siegel et al., 2016), and recent resting functional connectivity work suggests that interhemispheric disconnection is a significant predictor of apraxia severity (Watson et al., 2019). A combined SVR-CLSM and SVR-VLSM approach provides a rigorous analytic pipeline to further elucidate the degree to which inter- and intra-hemispheric disconnection predicts apraxia severity.

## Conclusion

For decades, the theoretical consensus has been the left inferior parietal lobule is the locus of manipulation knowledge, as lesions to this site are associated with apraxia. Our results offer a novel interpretation of the role of the left inferior parietal lobule in tool use: Tool manipulation knowledge is not “stored” local to the left inferior parietal lobule; rather, the left inferior parietal lobule is “hub-like” in function, as it aggregates **(i)** knowledge of action and object representations processed in ventral and lateral posterior temporal cortices with **(ii)** online visuo-motor mechanisms supporting the programming of hand and arm actions, and **(iii)** top-down feedback via frontal-motor structures to support the selection and implementation of tool use actions on the basis of task goals, rules, or context. Apraxia can arise from frank parietal damage, or due to damage of the white matter interfacing nodes of the Tool Use Network. It will be important for future studies to combine lesion-symptom mapping techniques with structural and functional connectivity measures to test causal hypotheses of the relative contributions to praxis made by distributed action and object representations in the human brain.

## Acknowledgements

We thank Cortney Howard, Leyla Tarhan, Louisa Smith, and Veronica Kreter for coding participants’ gestures, and Austin Wild, Olu Faseyitan, and Branch Coslett for help with lesion segmentation. Preparation of this manuscript was supported by a Moss Rehabilitation Research Institute/University of Pennsylvania postdoctoral training fellowship (NIH 5T32HD071844-05), by NIH grant R01 NS099061 to L.J.B., and by the General Electric-National Football League Head Health Challenge, Contract W911NF-09-0001 and Cooperative Agreement W911NF-19-2-0026 with the Army Research Office of the Army Research Laboratory to S.T.G.

## Author Information

The authors declare no competing financial interests. Correspondence should be addressed to F. Garcea.

## Footnotes

1 Importantly, there is involvement of the left inferior frontal gyrus in the uncorrected (*p* < .05) whole-brain disconnectome (see Supplemental Figure 3.

