## Supplemental Online Materials for "Structural Disconnection of the Tool Use Network After Left Hemisphere Stroke Predicts Limb Apraxia Severity"

**Supplemental Table 1.** Average MNI coordinate of each left hemisphere region in the Lausanne Atlas. Numbers next to each label denote subregion identity.

| Region Label | Center of Mass (Mean XYZ) | | |
|---|---|---|---|
| Left Lateral Orbitofrontal Cortex 1 | -28 | 23 | -18 |
| Left Lateral Orbitofrontal Cortex 2 | -19 | 43 | -19 |
| Left Pars Orbitalis | -43 | 39 | -13 |
| Left Frontal Pole | -8 | 67 | -10 |
| Left Medial Orbitofrontal Cortex | -7 | 39 | -18 |
| Left Pars Triangularis | -46 | 32 | 1 |
| Left Pars Opercularis | -48 | 17 | 14 |
| Left Rostral Middle Frontal Gyrus 1 | -36 | 33 | 32 |
| Left Rostral Middle Frontal Gyrus 2 | -32 | 49 | 20 |
| Left Rostral Middle Frontal Gyrus 3 | -30 | 59 | -3 |
| Left Superior Frontal Gyrus 1 | -9 | 56 | 13 |
| Left Superior Frontal Gyrus 2 | -8 | 44 | 36 |
| Left Superior Frontal Gyrus 3 | -17 | 25 | 52 |
| Left Superior Frontal Gyrus 4 | -10 | 3 | 61 |
| Left Caudal Middle Frontal Gyrus | -36 | 11 | 47 |
| Left Precentral Gyrus 1 | -17 | -21 | 67 |
| Left Precentral Gyrus 2 | -34 | -16 | 61 |
| Left Precentral Gyrus 3 | -46 | -6 | 43 |
| Left Precentral Gyrus 4 | -52 | 3 | 19 |
| Left Paracentral Gyrus | -7 | -29 | 59 |
| Left Rostral Anterior Cingulate Cortex | -6 | 38 | 1 |
| Left Caudal Anterior Cingulate Cortex | -5 | 20 | 27 |
| Left Posterior Cingulate Cortex | -5 | -18 | 38 |
| Left Isthmus Cingulate Cortex | -7 | -45 | 20 |
| Left Postcentral Gyrus 1 | -22 | -34 | 70 |
| Left Postcentral Gyrus 2 | -44 | -27 | 54 |
| Left Postcentral Gyrus 3 | -56 | -14 | 26 |
| Left Supramarginal Gyrus 1 | -54 | -28 | 24 |
| Left Supramarginal Gyrus 2 | -55 | -44 | 38 |
| Left Superior Parietal Lobule 1 | -30 | -46 | 60 |
| Left Superior Parietal Lobule 2 | -24 | -61 | 56 |
| Left Superior Parietal Lobule 3 | -16 | -81 | 39 |
| Left Inferior Parietal Lobule 1 | -42 | -77 | 22 |
| Left Inferior Parietal Lobule 2 | -40 | -65 | 40 |
| Left Precuneus 1 | -7 | -49 | 53 |

| | | | |
|---|---|---|---|
| Left Precuneus 2 | -9 | -64 | 30 |
| Left Cuneus | -5 | -81 | 18 |
| Left Pericalcarine Cortex | -11 | -82 | 5 |
| Left Lateral Occipital Cortex 1 | -18 | -98 | 6 |
| Left Lateral Occipital Cortex 2 | -38 | -83 | -1 |
| Left Lingual Gyrus 1 | -11 | -78 | -8 |
| Left Lingual Gyrus 2 | -17 | -57 | -2 |
| Left Fusiform Gyrus 1 | -37 | -65 | -14 |
| Left Fusiform Gyrus 2 | -34 | -27 | -28 |
| Left Parahippocampal Gyrus | -25 | -31 | -19 |
| Left Entorhinal Cortex | -25 | -5 | -33 |
| Left Temporal Pole | -29 | 13 | -38 |
| Left Inferior Temporal Gyrus 1 | -49 | -14 | -35 |
| Left Inferior Temporal Gyrus 2 | -54 | -53 | -12 |
| Left Middle Temporal Gyrus 1 | -61 | -47 | 0 |
| Left Middle Temporal Gyrus 2 | -56 | -8 | -23 |
| Left Bank of the Superior Temporal Sulcus | -53 | -46 | 8 |
| Left Superior Temporal Gyrus 1 | -57 | -37 | 14 |
| Left Superior Temporal Gyrus 2 | -52 | -2 | -12 |
| Left Transverse Temporal | -45 | -22 | 9 |
| Left Insula 1 | -37 | -12 | 5 |
| Left Insula 2 | -35 | 12 | -4 |
| Left Thalamus | -12 | -18 | 6 |
| Left Caudate | -13 | 8 | 10 |
| Left Putamen | -25 | 1 | -1 |
| Left Pallidum | -19 | -3 | -2 |
| Left Accumbens Area | -7 | 9 | -9 |
| Left Hippocampus | -25 | -23 | -14 |
| Left Amygdala | -24 | -3 | -21 |

**Supp. Fig. 1.** Edge-based Structural Disconnection Overlap among the 57 LCVA Participants.

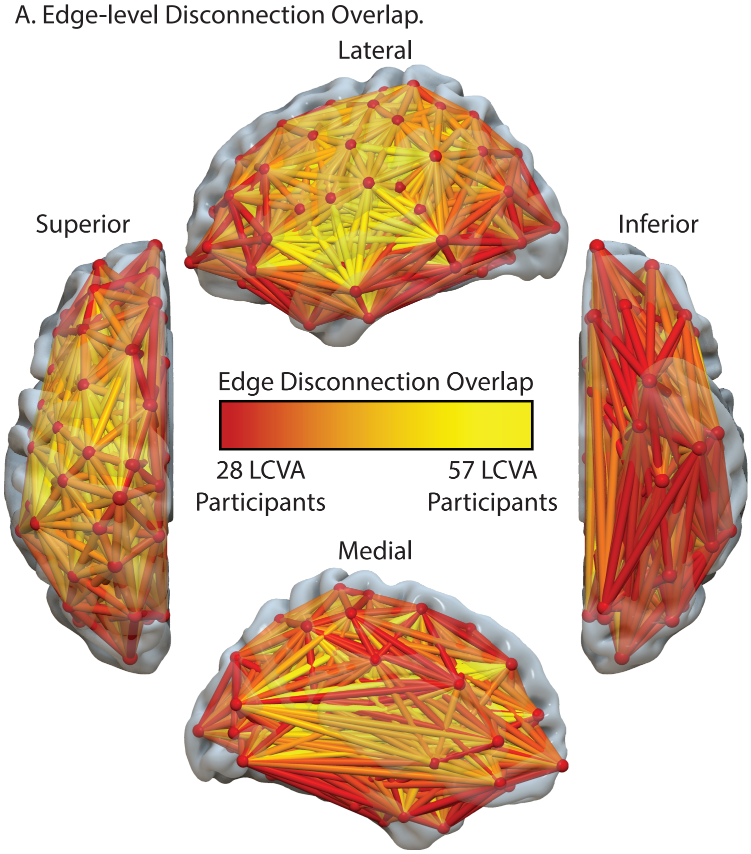


**Supplemental Figure 1. Edge-based Structural Disconnection Overlap among the 57 LCVA Participants. A.** Edge-based overlap among 57 participants. Only voxels with at least 28 participants are plotted. Note there is maximal disconnection throughout the left hemisphere.

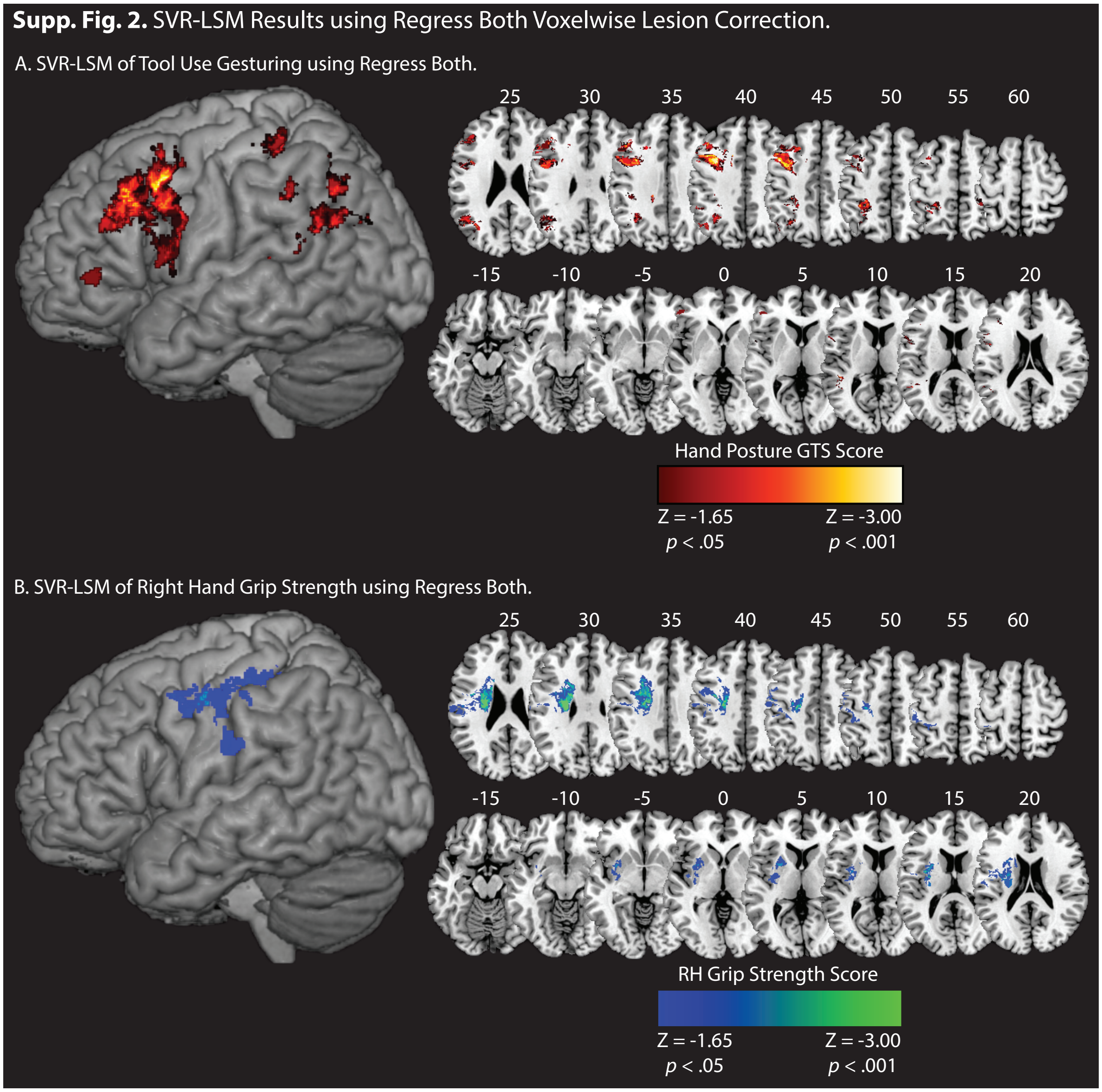


**Supplemental Figure 2. SVR-LSM of Hand Posture Errors in Gesturing Tool Use to the Sight of Objects. A.** Voxels associated with reduced performance in tool use gesturing (greater hand posture errors in tool use gesturing controlling for hand posture errors in meaningless imitation; red-to-white scale) using the 'Regress Both' lesion volume correction. We further removed significant voxels that did not form a cluster of at least 100 contiguous $1mm^3$ voxels. The analysis identifies two clusters. The first is in the left supramarginal gyrus and includes posterior superior temporal gyrus, the angular gyrus, and post-central gyrus. A second cluster is identified in the left middle frontal gyrus, and overlaps with the left inferior frontal gyrus (pars opercularis), pre-central gyrus, and the anterior portions of pars triangularis. **B.** Voxels associated with reduced right-hand grip strength (weaker grip strength with the right hand controlling for grip strength of the left hand; blue-to-green scale) identify the left pre- and post-central gyri, and subcortical regions including the insula, basal ganglia (caudate, putamen), portions of the frontal operculum, and the superior corona radiata. Whole-brain results are rendered in MNI space in 5-mm increments. SVR-LSM maps are set to a voxelwise threshold of $p < .05$ with 10,000 iteration Monte Carlo style permutation analysis.

**Supp. Fig. 3.** Left Hemisphere SVR-CLSM of Errors in Tool Use Gesturing.

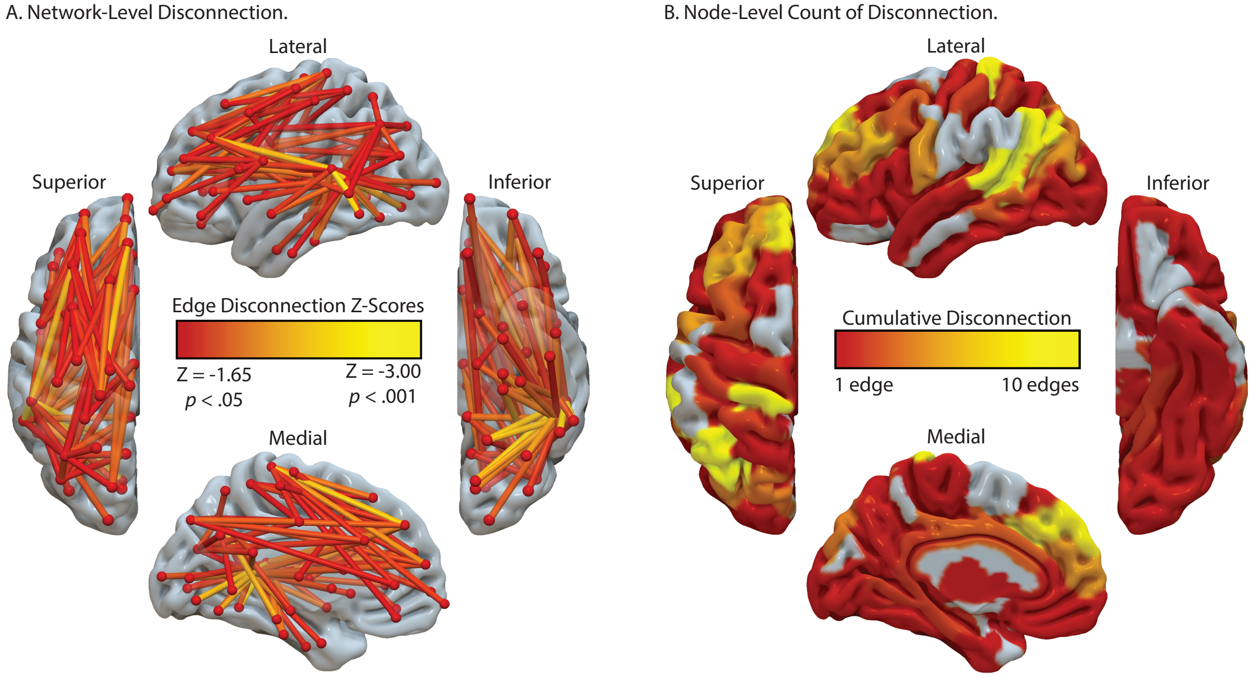


**Supplemental Figure 3. Left Hemisphere SVR-CLSM of Errors in Tool Use Gesturing. A.** The result of the SVR-CLSM identifies a disconnected fronto-temporal-parietal network associated with poor tool use gesturing (controlling for meaningless imitation). Note that these are the significant edges that were entered into the algorithm that identifies the maximally disconnected subgraph (see Figure 3). **B.** A node-level count of the disconnection identifies nodes in the left inferior parietal lobule, superior temporal gyrus, middle temporal gyrus, pre- and post-central gyri, and inferior, middle, and superior frontal gyri as exhibiting increased disconnection in association with poor tool use gesturing performance.

**Supp. Fig. 4.** Left Hemisphere SVR-CLSM of Reduced Right Hand Grip Strength.

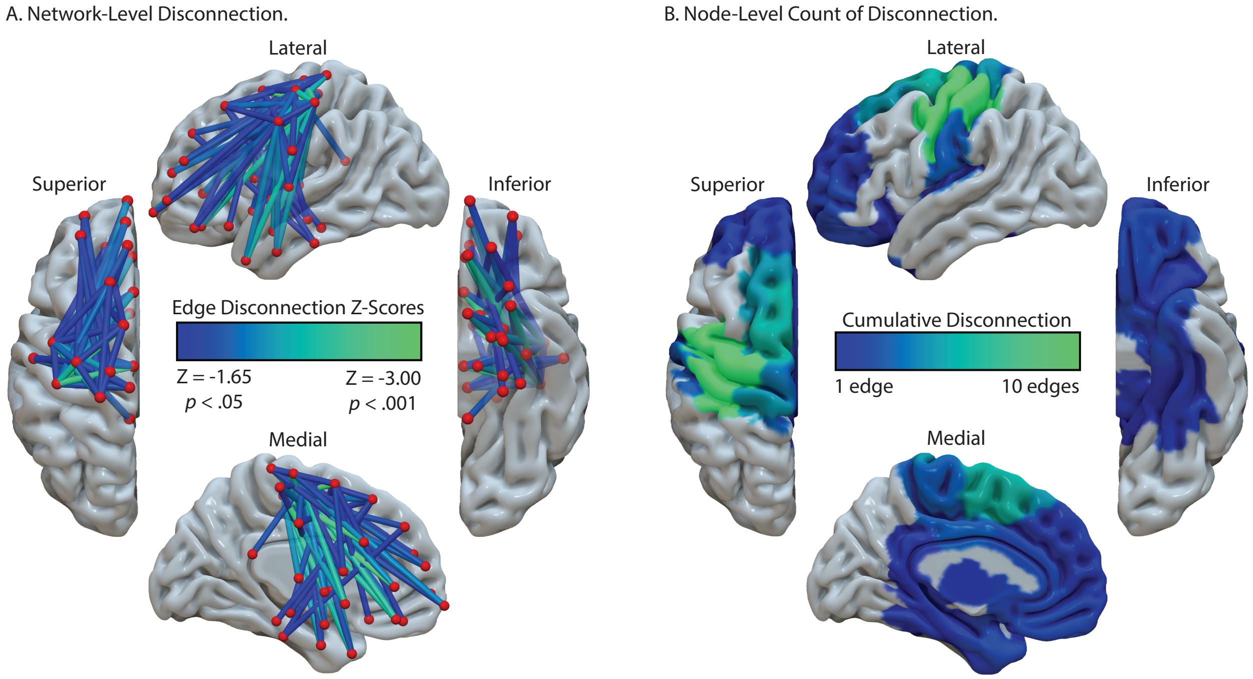


**Supplemental Figure 4. Left Hemisphere SVR-CLSM of Reduced Right-Hand Grip Strength. A.** The result of the SVR-CLSM identifies a disconnected fronto-motor network associated with of reduced right-hand grip strength (controlling for variability in left-hand grip strength). Note that these are the significant edges that were entered into the algorithm that identifies the maximally disconnected subgraph (see Figure 4). There is increased disconnection between pre- and post-central gyri and subcortical regions, consistent with the results of the SVR-LSM (Figure 2C). **B.** A node-level count of the disconnection identifies nodes in the pre- and post-central gyri, and superior frontal gyrus (~ dorsal premotor cortex) as exhibiting increased and widespread disconnection in association with reduced right-hand grip strength.

**Supp. Fig. 5.** Anatomical Proximity between the left posterior inferior parietal lobule and left SLF.

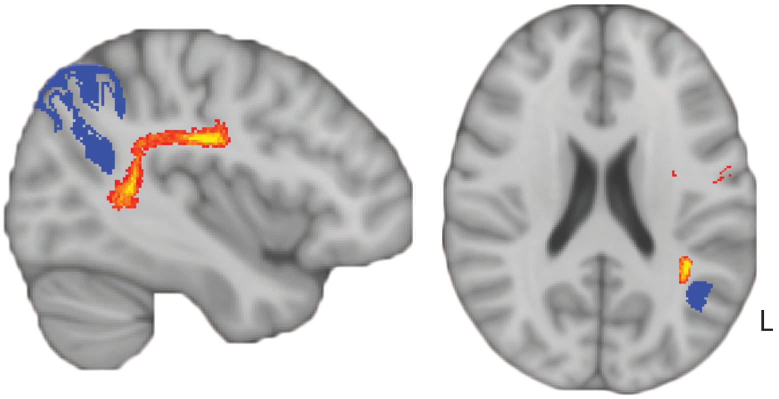


**Supplemental Figure 5. Anatomical proximity between the left posterior inferior parietal lobule and the left superior longitudinal fasciculus.** Voxels of the left posterior inferior parietal lobule (blue) are identified as structurally disconnected in association with reduced tool use gesturing performance and are situated in close anatomical proximity to the posterior portion of the left superior longitudinal fasciculus (SFL; red-to-yellow). The SLF is defined by the Johns Hopkins White Matter Atlas in FSL, set to a minimum probability value of 20% (red) and a maximum value of 50% (yellow).
